# LSD1 Performs Demethylase-Independent and Context-Specific Roles in Ewing Sarcoma

**DOI:** 10.64898/2025.12.04.692414

**Authors:** Rachel D. Dreher, Cenny Taslim, Ira Miller, John W. Sherman, Ariunaa Bayanjargal, Emily R. Theisen

**Affiliations:** Center for Childhood Cancer Research, The Abigail Wexner Research Institute at Nationwide Children’s Hospital, Columbus, OH, 43215, USA; Medical Scientist Training Program, College of Medicine, The Ohio State University, Columbus, OH, 43210, USA; Biomedical Sciences Graduate Program, College of Medicine, The Ohio State University, Columbus, OH, 43210, USA; Department of Pediatrics, College of Medicine, The Ohio State University, Columbus, OH, 43210, USA

**Keywords:** LSD1, Ewing sarcoma, nonenzymatic functions, transcriptomics, UM171, OG-L002

## Abstract

<u>L</u>ysine <u>s</u>pecific <u>d</u>emethylase 1 (LSD1), encoded by the gene *KDM1A*, is overexpressed and correlates with poor patient prognosis in Ewing sarcoma. LSD1 and the pathognomonic fusion oncoprotein, EWSR1::FLI1, colocalize throughout the genome, suggesting LSD1 is a critical co-regulator driving the progression of Ewing sarcoma. However, therapeutic targeting of LSD1 by competitive and noncompetitive inhibitors has yielded mixed results. Irreversible, enzymatic inhibition seems ineffective, but reversible noncompetitive inhibition has predominant off target mechanisms, leaving open the question of LSD1 function in Ewing sarcoma. Here we take a robust approach through multiple methods of depletion in multiple EwS cell lines to define enzymatic and nonenzymatic contributions of LSD1 to transcriptional regulation. We define a core set of 22 genes that are commonly repressed by LSD1 in all cell lines, and that repression of these genes downregulates synapse functioning and e-cadherin target genes. Derepression of these genes with LSD1 loss is an early and sustained genotype in all cell lines tested. We further define distinct gene sets in each cell line that are regulated by enzymatic and nonenzymatic LSD1 activity and find repression of e-cadherin target genes to be nonenzymatically regulated. This finding supports the growing body of evidence that in addition to their canonical catalytic activity, chromatin regulatory enzymes serve essential noncanonical roles as well. Furthermore, we uncovered evidence through use of the irreversible inhibitor OG-L002 that 2D cytotoxicity and proliferation assays may be insufficient to determine Ewing sarcoma response to LSD1 inhibition.

**SIGNIFICANCE:** Here we address a long-standing question in the field surrounding LSD1 and define the distinct enzymatic and nonenzymatic functions of LSD1 in Ewing sarcoma. In doing so, we have created a robust data set using genetic and pharmacological techniques in multiple models to thoroughly characterize LSD1 function in Ewing sarcoma cell lines.

## INTRODUCTION

Histone specific lysine demethylase 1 (LSD1) was the first chromatin demethylase identified, revealing histone methylation to be dynamically regulated^1^. LSD1 is comprised of 3 main structured domains; an N terminal SWIRM domain important for protein-protein interactions and substrate recognition, an elongated tower domain essential for key protein-protein interactions, and a C-terminal bi-lobed amine oxidase domain housing the catalytic pocket^2,3^. Through an FAD-dependent oxidative reaction, LSD1 removes mono- or di-methyl marks from lysine residues on the histone H3 tail resulting in either an open or closed chromatin state depending on which lysine residue is targeted^4^. Demethylation of histone 3 lysine 4 (H3K4) is repressive, and demethylation of lysine 9 (H3K9) is activating^5^. Despite the nomenclature, LSD1 can also demethylate non-histone proteins, including p53^6^, mef2d^7^, and HIF-1α^8^. LSD1 lacks a DNA binding domain and therefore must be incorporated into a larger protein complex to access its substrate in chromatin^9^. Most commonly, LSD1 binds to a SANT domain containing protein, such as CoREST 1/2/3 or MTA 1/2/3, within a larger chromatin regulatory complex (i.e. REST, BHC, NuRD)^10,11^ where it also serves a key structural role by recruiting histone deacetylases (HDACs)^12^. In concert with other complex members, LSD1 binds and demethylates the target lysine on the H3 tail shaping the local chromatin state.

The full scope of physiologic LSD1 functions has been reviewed elsewhere^13–15^. Since chromatin state plays an important role in development, LSD1 is critical for regulating the balance between stem cell pluripotency and lineage specific gene expression programs in early development. In embryonic stem cells, LSD1 promotes stem cell proliferation and expression of pluripotency genes^16–18^, but in other contexts, LSD1 induces skeletal muscle or osteogenic differentiation^7,19–21^ . In nonneuronal tissues, LSD1 represses neuronal specific genes to allow for non-neuronal differentiation^22^. Disruptions in differentiation and developmental state are hallmarks of cancer, and so aberrant LSD1 expression and activity have been implicated in numerous disease states as well. LSD1 is overexpressed in many high-grade sarcomas including Ewing sarcoma, rhabdomyosarcoma, and osteosarcoma^23,24^. Ewing sarcoma (EwS) cell lines have the second highest levels of LSD1 RNA overexpression behind neural cancers, and higher levels of LSD1 correlate with poor patient prognosis^25^, making LSD1 an attractive therapeutic target. EwS is a devastating adolescent and young adult cancer, with no targeted or precision treatments available^26,27^. In 85% of cases, EwS is driven by the t(11;22)(q24;q12) chromosomal translocation, resulting in the fusion oncogene *EWSR1::FLI1*^28^. The remainder of cases are driven by alternative FET::ETS fusions, such as *EWSR1::ERG*^27^. Unfortunately, these fusion transcription factors remain undruggable due to the lack of a concave binding pocket and the disordered FET domain. In response, extensive work has been done to determine whether targeting coregulators like LSD1 could disrupt the downstream oncogenic activity of the fusion. LSD1 is overexpressed, correlates with prognosis, and colocalizes with the fusion in multiple cell lines, supporting LSD1 as a promising therapeutic vulnerability^25,29^. As such, LSD1 inhibition has been pursued, but results have been difficult to interpret. Irreversible enzymatic inhibition of LSD1 via the tranylcypromine (TCP) derivative GSK-LSD1, seemingly has no cytotoxic effect on EwS cell lines^25^. In contrast, small molecule noncompetitive inhibition of LSD1, targeting enzymatic and nonenzymatic functions, is potently cytotoxic in EwS cell lines, but we have recently shown this inhibitor is predominantly acting through off target mechanisms^30^. Despite considerable interest in LSD1 as a therapeutic target in EwS, disparate results with different inhibitors pose challenges for interpreting its function and relevance yet highlights the possibility of significant nonenzymatic functions of LSD1.

The specific nonenzymatic functions of LSD1 in Ewing sarcoma have yet to be defined. LSD1 has mostly been studied for its enzymatic, demethylase function, but there is a growing body of evidence that it has nonenzymatic functions, such as scaffolding, in other contexts. Within the CoREST complex, LSD1 is required for recruitment and proper functioning of HDACs^12,31^. Other DNA binding proteins such as GFI1B, ZNF217, and SNAI1 recruit larger repressive complexes through binding LSD1^3,18,32,33^. LSD1 scaffolding also stabilizes DNMT1 to preserve the DNA methylation signature essential for proper ESC differentiation^34^. Based on these findings, we hypothesize that LSD1 has distinct nonenzymatic roles in EwS. We have created a molecular suite of tools to deplete LSD1 via multiple methods to define the distinct set of genes regulated by LSD1 and to delineate the enzymatic and nonenzymatic functions of LSD1 in EwS.

## MATERIALS AND METHODS

### Reagents and Constructs

OG-L002 was purchased from Selleck Chemicals (Cat. # S7237). UM171 was purchased from MedChemExpress (Cat# HY-12878). shRNA constructs against luciferase (iLuc) and LSD1 (iLSD1) as well as pMSCV_empty vector (puro) have been previously described^25,35^. HA- and twinstrep-tagged LSD1wt and LSD1ed were generated in house. pMCSV_puro was first digested with XhoI as described by NEBcloner. Wild type LSD1 sequence was PCR amplified out of a preexisting plasmid (pET28a_hLSD1) and a gene block with the twinstrep and 3xHA tag was ordered from Genscript. These two sequences were cloned into the pMSCV_puro backbone following Gibson cloning ratios. LSD1ed was generated from the LSD1wt plasmid by digesting with BamH-HF followed by BsaBI with heat inactivation. A Genscript gene block with the K661Q and A539E mutation was then cloned into the digested LSD1wt back bone following Gibson cloning ratios. All plasmid sequences were verified by sequencing (Plasmidsaurus). All gene block sequences, PCR primers are included in Supplementary Table 1.

### Cell Lines and Tissue Culture

All Ewing sarcoma cell lines A673 (RRID: CVCL_0080), CHP-100 (RRID: CVCL_7166), TTC-466 (RRID: CVCL_A444) were originally provided by Dr. Stephen Lessnick. A673 cells were cultured in Dulbecco’s Modified Eagle Medium (DMEM; Corning Cellgro 10-013-CV) supplemented with 10% fetal bovine serum (FBS: Gibco 16000-044), 1% penicillin/streptomycin/glutamine (P/S/Q; Gibco 10378-016), and 1% sodium pyruvate (Gibco 11360-070). CHP-100 cell lines were maintained in McCoys 5A Medium (Gibco, 16600082) supplemented with 10% heat inactivated fetal bovine serum (heated for 30 min at 56°C), and 1% PSQ. TTC-466 cell lines were cultured in Roswell Park Memorial Institute 1640 Medium (RPMI 1640; Corning Cellgro 15-040-CV) supplemented with 10% FBS and 1% PSQ. All cells were maintained in an incubator at 37°C and 5% CO_2_. Every experiment was performed within 2 months of thawing a cell line and all cell lines were tested for mycoplasma annually using the Universal Mycoplasma Detection Kit (ATCC 30-1012K) and STR profiled (LabCorp) upon thawing.

### LSD1 Knockout Line Generation

LSD knock out (KO) cell lines were created from A673, CHP-100, and TTC-466 parental cell lines, in collaboration with the Nationwide Children’s Hospital Abigail Wexner Research Institute’s CRISPR core. The core designed CRISPR guide RNAs against *KDM1A* and used an electroporation delivery protocol optimized for Ewing sarcoma cell lines to deliver the guides preloaded onto the Cas9 enzyme. Guides targeted exon 17 and 18 of the LSD1 gene to prevent transcription of a fully functional mature mRNA strand. Guide sequences are listed in Supplementary Table 1. Surviving cells were allowed to grow until confluent, then the pellet was tested for LSD1 KO via sequencing and immunoblotting. All KO cell lines are stable and KO in the polyclonal population was confirmed by immunoblot with every experiment.

### Antibodies

The antibodies used for immunodetection are as follows; anti-LDS1 (Cell Signaling C69G12, RRID:AB_2799915), anti-lamin (Abcam ab133741, RRID:AB_2616597), anti-HA (Abcam ab18181, RRID:AB_444303), IRDye 800CW goat anti-mouse IgG (LI-COR Biosciences, 926-32210, RRID: AB_621842), and IRDye 800CW goat anti-rabbit IgG (LI-COR Biosciences, 926-32211, RRID: AB_621843).

### Retrovirus Production and Transduction of Cells

All shRNA and rescue constructs were delivered to cells via retroviral transduction. Retrovirus was produced using HEK-239T-EBNA cells stably expressing Ebstein-Barr nuclear antigen (EBNA) under neomycin selection. 10µg of cDNA vector, vsv-g, and gag-pol were delivered in separate plasmids using Optimem and Mirus to transfect the HEK-293T-EBNA cells. After two days, virus was collected in viral collection media (DMEM with 10% heat inactivated FBS, 1% P/S/Q, 2% 1M HEPES pH 7.5) in aliquots over two days. Virus was passed through a 2µM filter and immediately added to cells to be infected. Target cells were transduced with retrovirus supplemented with 2.5 µL of 8 mg/mL polybrene. Cells incubated in the viral mix for 2 hours then were supplemented with 8mL of fresh media and 8 µL of 8 mg/mL polybrene. The next day cells were passed into selection media containing 2 µg/µL puromycin (Sigma P8833) for selection for 3 days. Following selection cells were seeded for Incucyte cell proliferation assays and soft agar assays, and pellets were collected for protein and RNA isolation for subsequent western blot analysis and RNA extraction for RNA-sequencing.

### Incucyte Cell Proliferation Assay

Ewing sarcoma cells were seeded at a density of 2,000 cells per well for A673 and CHP-100 and 5,000 cells per well for TTC-466 in triplicate in a 96 well clear tissue culture plate (Corning). Phase Contrast images were taken every 4 hours until confluent or growth arrest (approximately 9 days) in the Incucyte ZOOM Kinetic Imaging System (Essen BioScience). Cell confluence was measured using Incucyte ZOOM 2016A software (Essen BioScience) and plotted to display a growth curve. Each data point represents the mean +/- standard error of the mean (SEM) from a minimum of 3 replicates.

### Soft Agar Colony Formation Assay

Soft agar assays were performed as previously described^25,36^. For A673 and CHP-100 cell lines, 7,500 cells were seeded per agar, and for TTC-466 18,000 cells were seeded per agar. Every plate was fed twice a week with 500 µL of media supplemented with selection antibiotic or drug and then imaged at 2 weeks post seed. Colonies were counted at 2 weeks using ImageJ software.

### Dose Response Curves

10,000 cells per well in 200 µL of media were plated in triplicate in a 96 well clear tissue culture plate (Corning). After 24 hours, a 9-part serial dilution of UM171 in media was prepared in PCR tubes. Equivalent volumes of the drug dilution were added to each of the wells. Following 96 hrs of incubation, 110µL of media was removed from each well and replaced with 80µL of CellTiter-Glo (Promega) reagent. The plate was covered with aluminum foil to reduce light and agitated at 300 rpm for 15 min at room temperature. Afterwards, 160µL of the cell solution was transferred to an opaque walled 96 well plate (Corning) and luminescence was measured on a Synergy H1 plate reader (Agilent). Cell viability was calculated relative to an untreated DMSO control well.

### Protein Isolation and Western Blot Analysis

1-5 million cells per sample were pelleted then flash frozen at the desired time point post drug treatment of viral infection. Whole cell protein was extracted using Pierce RIPA buffer and the supernatant was transferred to a new tube for quantification by Pierce^TM^ BCA Protein Assay Kit (Thermo Scientific 23225). 10 µg of each protein sample was run on a precast 4-15% gradient Tris-Glycine gel (BioRad 4561084) at 90V for 20 min then 110V for 1 hour. Proteins were then transferred onto a nitrocellulose membrane (Thermo IB23002) using a Trans-Blot SD Semi-Dry Transfer Cell (Bio-Rad, 1703940) at 15V for 1 hour. The membranes were then blocked for 60 minutes at room temperature with Odyssey® RBA Blocking Buffer (Li-Cor 927-40003). Primary antibody was applied at 1:1000 concentration overnight at 4°C. The next day, the membrane was washed 3 times for 5 minutes with TBST (Tris Buffered Saline with Tween 20), then goat anti-rabbit IgG secondary antibody was applied for 1 hour at room temperature. Membranes were washed as previously described and imaged on the Odyssey® LI-COR CLx Imager (LI-COR Bio, discontinued).

### Quantitative reverse-transcriptase polymerase chain reaction (qRT-PCR)

Cell pellets of 200,000-500,000 cells were flash frozen with liquid nitrogen prior to preparation. Samples were thawed on ice and total RNA was extracted using an RNAeasy kit (Qiagen). RNA concentration was then measured on a NanoDrop (Thermo). Total RNA was amplified using one-step qRT-PCR and SYBR green fluorescence for quantification. Post amplification, a melting curve analysis was performed. Normalized fold enrichment was calculated relative to RPL30. Primer sequences are available in Supplementary Table 1.

### RNA Isolation for RNA Sequencing

250,000 cells were pelleted and flash frozen post treatment for RNA isolation and sequencing. Total mRNA was extracted from the cell pellets following the Qiagen RNeasy kit. A gDNA removal column and DNase treatment were used on all samples to purify for RNA. RNA concentrations were measured using a Nanodrop and 1µg of RNA per sample (20 µL of 50ng/µL) were submitted to the Abigail Wexner Research Institute Genomic Services Laboratory for library preparation (TruSeq Stranded mRNA Kit; Illumina 20020594), library quality control, and next generation sequencing (Illumina NovaSeq 6000). The target sequence depth was 50 million 150-bp paired end reads. The raw BCL files were converted to FASTQ files and returned for in house analysis.

### RNA Sequencing Analysis

FASTQ files from AWRI GSL were analyzed using an in house developed RNA sequencing analysis pipeline^37^. The resulting DESeq2 output was used for data visualization. Overlap analysis of commonly differentially expressed genes was performed using base R packages (DESeq2, ggplot2 (RRID: SCR_014601), VennDiagram, grid, pbapply). Pathway analysis was performed using the R package enrichR^38^ and clusterProfiler^39^. Gene set enrichment analysis was performed using GSEA_4.4.0 (GSEA; RRID: SCR_003199, https://www.gsea-msigdb.org/gsea/index.jsp). The direct A673 and TTC-466 fusion target list used has been previously defined^35,40^.

### Data Availability

The data sets analyzed in this work are deposited in the NCBI’s Gene Expression Omnibus (GEO) and Sequencing Read Archive and are accessible through GEO Series Accession GSE311851. All other data are available within the main manuscript, supplemental files, or upon request from the corresponding author.

## RESULTS

### LSD1 represses neural synaptic functioning and e-cadherin target genes in Ewing sarcoma

Ewing sarcoma (EwS) is a heterogenous disease, therefore we chose to study LSD1 functions using orthogonal methods in multiple cell lines. Our three-cell line panel, consisting of A673, CHP-100, and TTC-466, is derived from both primary tumor and metastases, includes 3 different fusions, and harbors both wild type and mutated versions of two of the most frequent patient mutations in Ewing sarcoma, p53 and CDKN2A^41^ (Table 1).

**Table 1:**
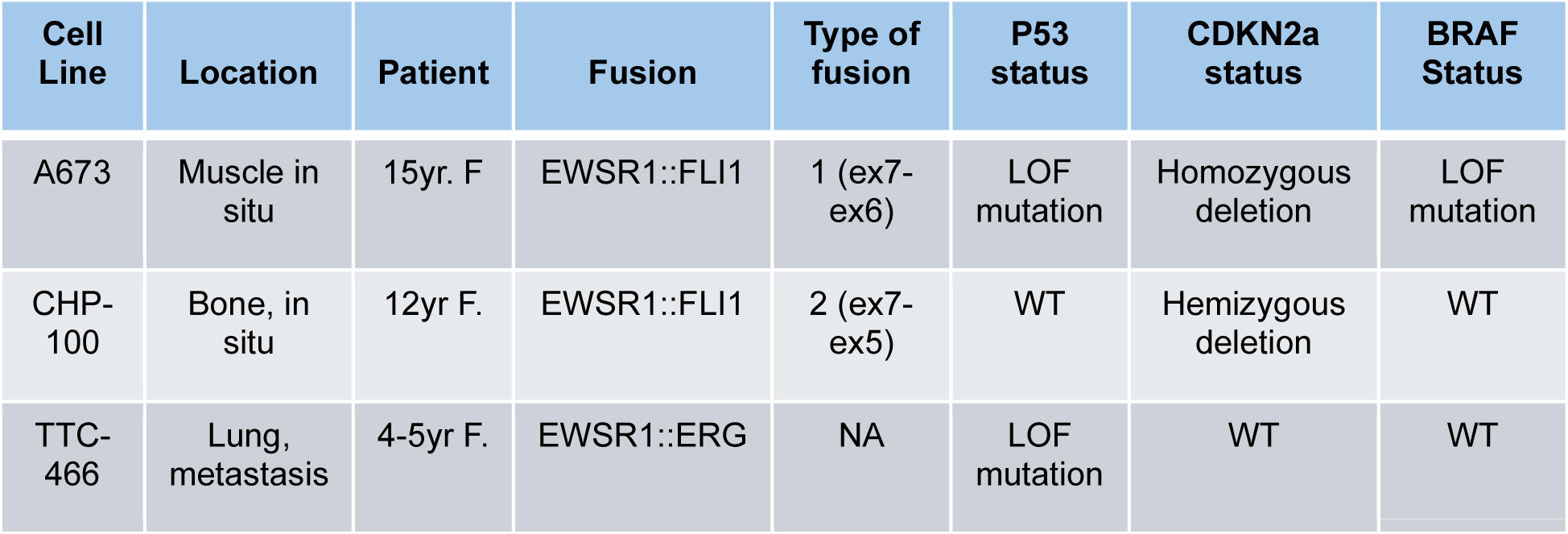
Ewing sarcoma cell line characteristics.

To define LSD1 function, we sought to deplete LSD1 at the DNA, mRNA, and protein level. We used CRISPR-Cas9 mediated gene editing to create unselected, stable, polyclonal LSD1 knockout (KO) cell lines from all three parental EwS lines. We also dosed parental EwS cells with 800nM of UM171 for 96 hrs to degrade the LSD1 protein (UM) (Figure 1A). UM171 functions as a molecular glue, stabilizing the interaction between its substrate and the KBTBD4 subunit of an E3 ligase allowing for ubiquitination and subsequent degradation^42,43^. In all three cell lines we achieved knockout with transient CRISPR-Cas9 editing and significant depletion of LSD1 protein with UM171(Figure 1B-D, Supplementary Figure 1A-B).

**Figure 1:**
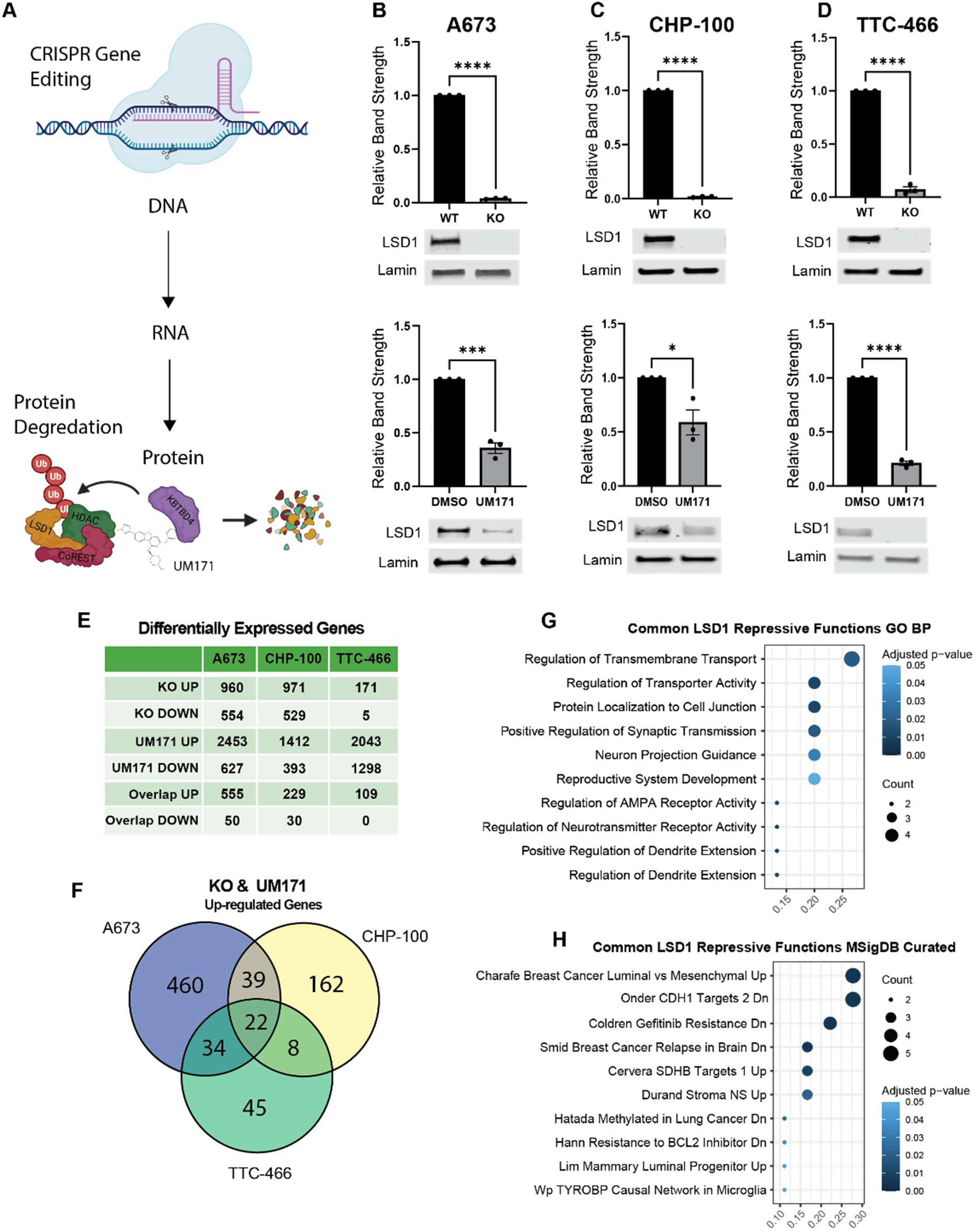
LSD1 represses synaptic functioning and CDH1 target genes. **(A)** Schematic of LSD1 depletion techniques targeting DNA via CRISPR Cas9 gene editing or protein via UM171 mediated degradation. For UM171 knockdown, parental cells were treated with DMSO or 800nM of UM171 for 96 hours. Schematic was created in BioRender. Representative western blot and densitometry quantification of triplicate samples of LSD1 KO and LSD1 knockdown in A673 **(B)** CHP-100 **(C)** and TTC-466 **(D)**. **(E)** Table of up- and down-regulated genes in all three cell lines of KO vs WT and UM171 vs DMSO. Overlap up and overlap down are the genes commonly up- or down-regulated between KO and UM171 in each cell line. **(F)** Venn diagram representation of genes commonly upregulated between KO and UM171 in all three cell lines. DEGs were defined as FDR > 0.05 and fold change > 2. Pathway enrichment analysis of core LSD1 target genes ranked by gene ratio from the **(G)** Gene Ontology Biological Processes (GOBP) database and the **(H)** MSigDB Curated database.

LSD1 knockdown via shRNA in A673 cells has been previously reported^25,44^. However, we were unable to get comparable knockdown across all cell lines using a consistent shRNA design (Supplementary Figure 1C). Furthermore, despite no knockdown in CHP-100 or TTC-466, we saw a modest reduction in 2D cellular proliferation, and a pronounced reduction in 3D growth in soft agar (Supplementary Figure 1D-F). Confounding factors thus hampered our ability to determine the effect of LSD1 loss with shRNAs. We proceeded using our consistently successful depletion methods, KO and UM for further analysis (Supplementary Figure 1G).

To identify transcriptional changes resulting from LSD1 loss, we performed RNA sequencing on our KO and UM samples. When analyzed together, cell line identity was the primary source of variation between samples, supporting the heterogenous nature of EwS (Supplementary Figure 1H). The number of differentially expressed genes (DEGs) varied greatly with mechanism of LSD1 depletion and cell line, with substantially more upregulated DEGs than downregulated DEGs consistent with LSD1’s defined repressive role (Figure 1E). Notably, LSD1 KO in TTC-466 only resulted in 176 total DEGs.

Despite the variability, we identified a core set of 22 genes that are commonly upregulated following KO or UM171 treatment across all cell lines (Figure 1F). There are no genes commonly down regulated across cell lines and conditions. We define these 22 upregulated genes as high confidence LSD1 regulated genes in EwS. Pathway analysis of these genes indicates LSD1 represses synaptic and neurotransmitter receptor activity, as well as e-cadherin (CDH1) target genes (Figure 1G-H). The pathway titled “Onder CDH1 Targets Dn” refers to genes that are downregulated with loss of CDH1 and are therefore activated by endogenous CDH1. Hence, LSD1 represses genes that are activated by CDH1.

Loss of CDH1 is a hallmark of epithelial to mesenchymal transition (EMT) and metastasis in tumor cells^45,46^. Repression of CDH1 targets therefore suggests LSD1 contributes to a pro-metastatic environment. Cell line specific pathway analysis reveals these common pathways are not always the most enriched following LSD1 depletion, but repression of CHD1 target genes consistently ranks among the top 2 enriched pathways by gene ratio in all cell lines (Supplementary Figure 3A-F).

We checked whether these core LSD1 genes are co-regulated by the oncogenic driver fusion. EWSR1::FLI1 and EWSR1::ERG are known to regulate distinct sets of genes in different cell lines^40^. We thus reanalyzed published RNA sequencing data from knockdown(KD)/rescue experiments in A673 and TTC-466 cells and generated new KD/rescue datasets for CHP-100 cells to define fusion regulated target genes in each cell line (Supplementary Figure 4J-K). We then defined fusion target genes as those commonly regulated in all 3 cell lines (Supplementary Figure 4A-B). Only 3 genes were present in the overlap between common LSD1 and common fusion genes, but we noted significant overlap in cell line-specific fusion gene targets, both activated and repressed, and LSD1 repressed targets (Supplementary Figure 4D-F). Despite significant overlap, gene set enrichment between LSD1 target genes and fusion targets were not significant in any cell line, indicating little to no functional relationship between LSD1 and fusion function in these cell lines (Supplementary Figure 4G-I).

### De-repression of e-cadherin target genes is early and sustained

The KO cell lines here are stable polyclonal populations, as cells have adapted following electroporation CRISPR editing and gene loss. In contrast, UM171-induced protein degradation allowed for temporal analysis of the changes in gene expression following loss of LSD1 protein. Six hours of UM171 treatment is the earliest time point we see sufficient LSD1 degradation (Supplementary Figure 5A-B). We therefore dosed our three cell lines with 800nM of UM171 for 6 hrs, 24 hrs, and 96 hrs to deplete LSD1 and analyzed each time point by RNA sequencing to identify early (direct) and late (indirect) gene targets of LSD1 (Figure 2A-C).

**Figure 2:**
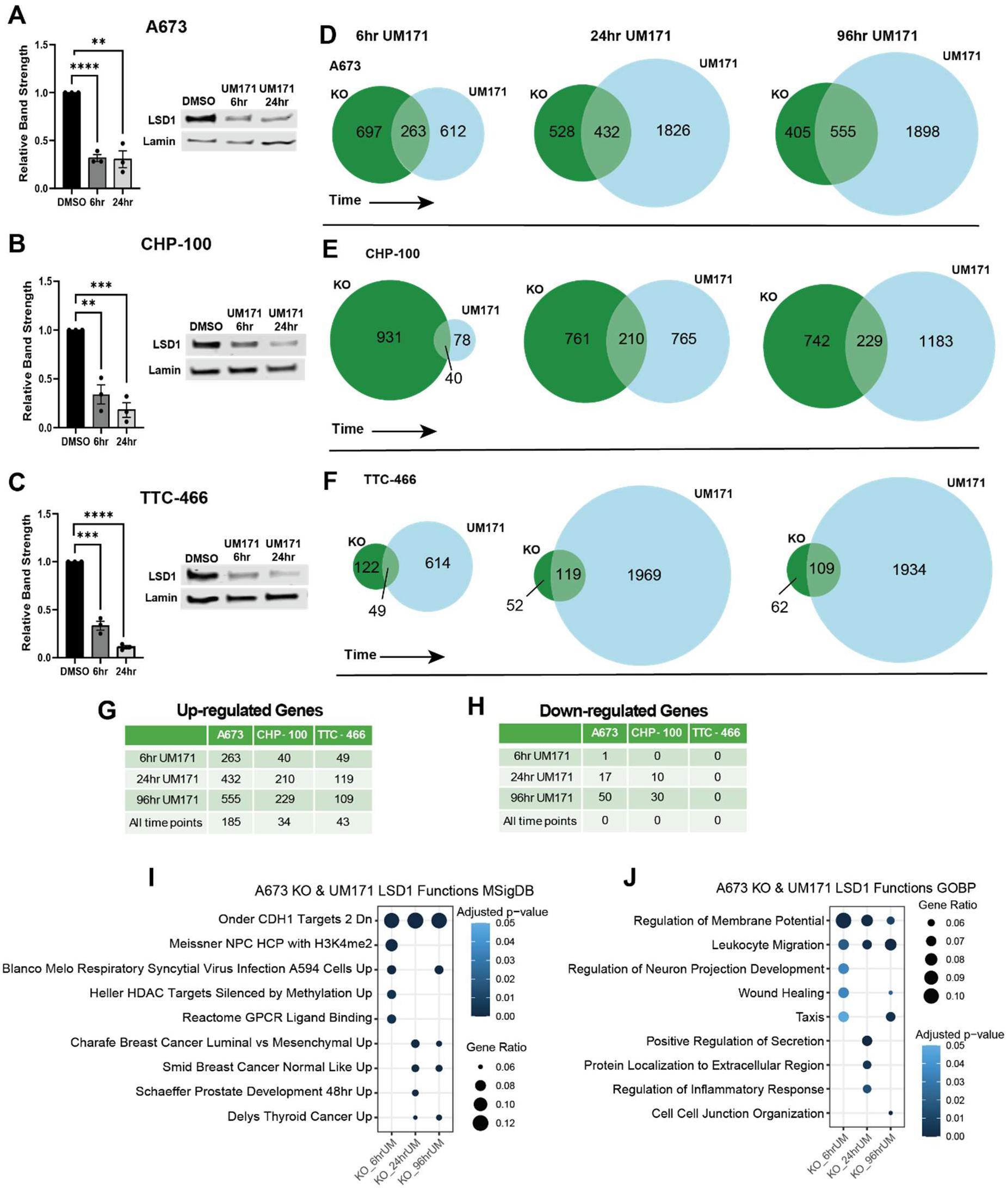
Derepression of CDH1 target genes is early and sustained post LSD1 degradation. **(A)** A673 **(B)** CHP-100 and **(C)** TTC-466 representative western blots and densitometry quantification of LSD1 degradation after DMSO and 800nM of UM171 for 6 hrs and 24 hrs. **(D-F)** Overlaps of genes commonly up-regulated between KO and UM171 at 6 hrs, 24 hrs, and 96 hrs in all three cell lines. A table summarizing genes commonly **(G)** up- and **(H)** down-regulated between KO and UM171 at 6 hrs, 24 hrs, 96 hrs, and all three time points in each of the three cell lines. Representative pathway analysis of A673 ranked by gene ratio between KO and UM171 at 6 hrs, 24 hrs, and 96 hrs using **(I)** MSigDB Curated and **(J)** GO BP databases.

We defined DEGs at each time point as those common to KO and UM171 treatment to minimize inclusion of non-LSD1 targets of UM171. As time progresses there are more overlapping upregulated genes between the KO and UM samples in all cell lines, and common between the three cell lines, likely corresponding to progressive downstream effects of LSD1 loss (Figure 2D-F, Supplementary Figure 5F-H). Interestingly, the number of genes regulated by LSD1 is strongly dependent on cell line, supporting a context-dependent role of LSD1 within EwS (Figure 2G-H, Supplementary Figure 5C-E).

Having identified LSD1’s common role in repressing CDH1 target genes across all cell lines, we performed pathway analysis on each time point in each cell line to determine the kinetics of derepression following LSD1 loss. Derepression of CDH1 target genes is apparent at 6 hrs of LSD1 depletion and sustained throughout 96 hrs of depletion in all cell lines (Figure 2I, Supplementary Figure 6A-D). This was surprising considering only 4 genes were common between all cell lines as early as 6 hrs of UM171 treatment (Supplementary Figure 5G). LSD1 likely performs this common function through slightly differing gene targets in each cell line.

### LSD1 degradation results in a cell line specific deficit in 3D proliferation in agar

Prior work has demonstrated significant impairment of 2D cellular proliferation and 3D growth in soft agar with shRNA-mediated LSD1 loss^25^. However as discussed above, shRNA-mediated depletion is inconsistent across cell lines. Therefore, we sought to determine whether loss of LSD1 via KO or UM171 degradation impacts 2D and 3D cellular proliferation as with shRNA knockdown and treatment with noncompetitive LSD1 inhibitors.

We were surprised that KO of LSD1 in two of three cell lines did not reduce cellular proliferation, with only a slight lag in proliferation in TTC-466 (Figure 3A-C, top panel). Furthermore, when KO cells are plated in soft agar, there is no reduction in 3D cellular proliferation in any cell line (Figure 3A-C, bottom panel). In TTC-466 the colonies appear smaller, but the number is no different than DMSO treatment.

**Figure 3:**
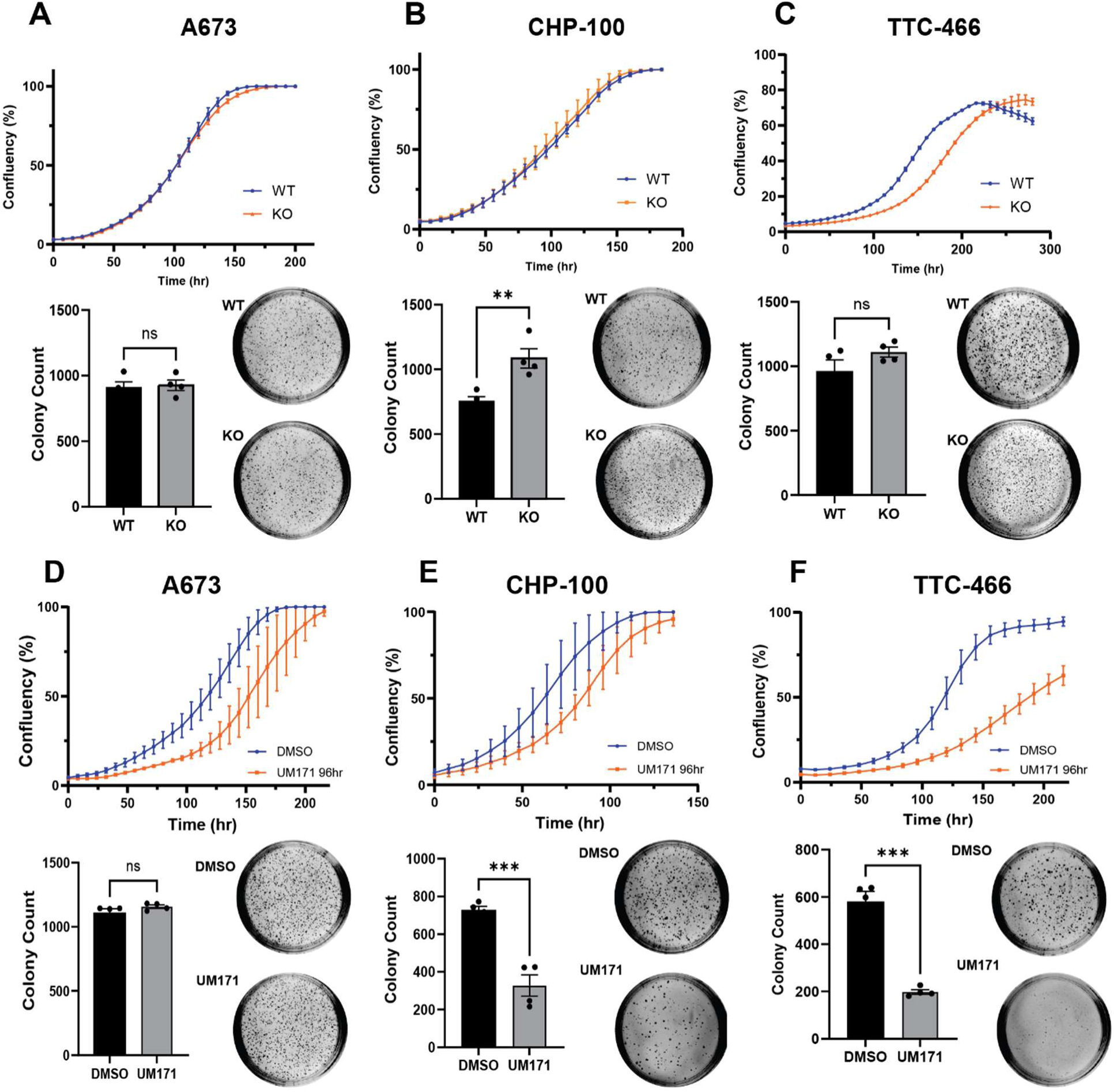
Loss of LSD1 results in cell line dependent defects in 3D proliferation in agar. (A-C) LSD1 KO vs WT 2D cellular proliferation was measured over a week in parallel with LSD1 KO vs WT 3D proliferation in soft agar in **(A)** A673, **(B)** CHP-100, and **(C)** TTC-466. Representative images of colonies in agar are shown with accompanying quantification of colonies across replicates. 2D and 3D proliferation were measure as above in DMSO vs 800nM of UM171 for 96 hrs in **(D)** A673, **(E)** CHP-100, and **(F)** TTC-466. Cells were pre-treated with DMSO or UM171 for 96 hrs prior to plating for assays.

We then measured cellular proliferation and 3D growth in soft agar with cells pre-treated for either 6 hrs or 96 hrs with UM171 and again saw a cell line specific response. A673 and CHP-100 cells showed a slight delay in 2D cellular proliferation, while TTC-466 has a more pronounced response (Figure 3D-F, top panel, Supplementary Figure 7A-C, top panel). We were again surprised to find different responses to treatment in anchorage independent growth in soft agar. Like KO, UM171 treatment did not reduce A673 cell growth in soft agar, but surprisingly in CHP-100 and TTC-466 we see a significant decrease in colonies when seeded after either 6 hrs or 96 hrs of UM171 treatment (Figure 3D-F, bottom panel,

Supplementary Figure 7A-C, bottom panel). This is not due to differing IC50s of UM171 between the cell lines or to differing levels of baseline LSD1 expression (Supplementary Figure 7D-F).

Stable KO of LSD1 does not create a phenotype, but early loss of LSD1 by degradation decreased 3D proliferation in a subset of cell lines. The differences in colony formation could be due to selective pressure exerted on cells during electroporation at genome editing, off target effects of UM171, or both. We therefore sought an alternative approach to test whether the effects seen in soft agar were related to impaired LSD1 function. To do this, we used an irreversible LSD1 inhibitor to directly and specifically block LSD1 demethylase function in EwS cells^30,47^.

### Enzymatic inhibition of LSD1 is cell line specific

OG-L002 is a tranylcypromine (TCP) derivative that blocks LSD1 demethylase activity by forming an adduct with the FAD cofactor (Figure 4A)^47^. Both OG-L002 and another TCP derivative, GSK-LSD1, showed minimal cytotoxicity in 2D assays of EwS cell growth, but 3D growth in agar following OG-L002 or GSK-LSD1 treatment has only been studied in A673 cells^25,48^. As expected, in A673 OG-L002 does not delay 2D proliferation, nor impair 3D colony growth in soft agar (Figure 4B). However, despite minimal defects in 2D proliferation in CHP-100 and TTC-466, we were surprised to find both cell lines showed a statistically significant reduction in colonies following OG-L002 treatment (Figure 4C-D). This finding suggests inhibiting LSD1 enzymatic activity could be more important for anchorage independent growth than proliferation, with cell line specific sensitivity to this treatment.

**Figure 4:**
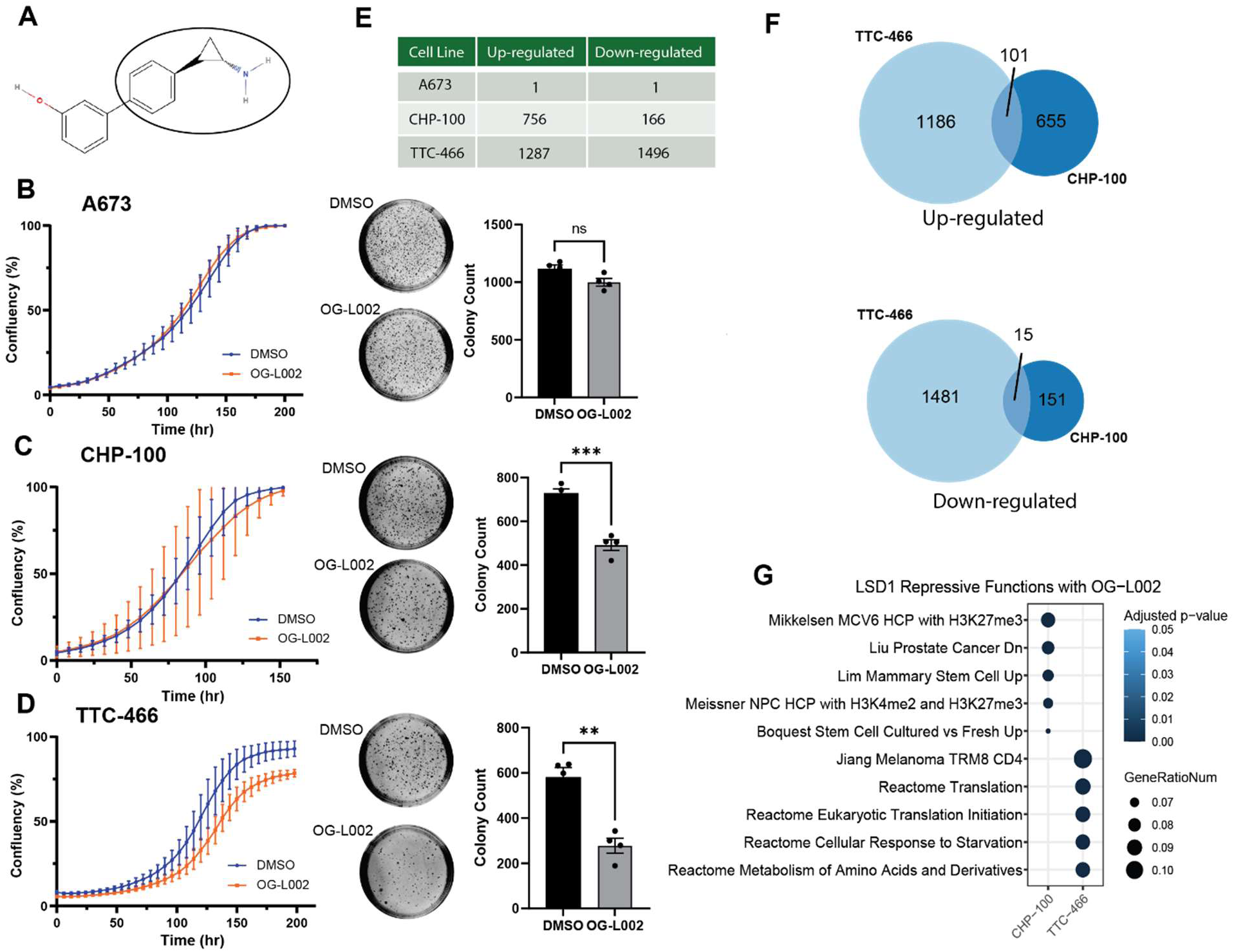
Enzymatic inhibition of LSD1 is cell line specific. **(A)** Structure of LSD1 enzymatic inhibitor OG-L002 with the moiety that forms the adduct with FAD circled. **(B-D)** 2D cellular proliferation of cells treated with DMSO or 2µM of OG-L002 for 96 hrs measured over 1 week. In parallel, 3D proliferation in agar of cells treated with DMSO or 2µM of OG-L002 for 96 hrs were measured over 2 weeks. Representative images of colonies in agar are shown with accompanying quantification of colonies across replicates in **(B)** A673, **(C)** CHP-100, and **(D)** TTC-466. **(E)** Table of cells up- and downregulated by OG-L002 treatment in each cell line. **(F)** Venn diagram representation of overlapping up- and downregulated genes common between CHP-100 and TTC-466. **(G)** Top 5 enriched pathways ranked by gene ratio from the MSigDB Curated database in CHP-100 and TTC-466 with OG-L002 treatment.

We performed RNA sequencing on our OG-L002 treated samples and found the degree of transcriptomic changes correlates with the cell line dependent reduction in 3D growth in soft agar (Figure 4E, Supplementary Figure 8A-B). There is only 1 upregulated DEG and 1 downregulated DEG in A673 with OG-L002 treatment, compared to 756 up- and 166 downregulated DEGs in CHP-100 and 1287 up-and1496 downregulated DEGs in TTC-466 (Figure 4E). There are no genes commonly regulated in all three cell lines, however 101 up- and 15 down-regulated DEGs are common between CHP-100 and TTC-466 (Figure 5F). The substantial difference between the number of DEGs and the few overlaps between CHP-100 and TTC-466 suggest LSD1 may have different enzymatic functions in each cell line. Therefore, we performed individual pathway analysis of each cell line to define the top 5 enriched pathways in CHP-100 and TTC-466 (Figure 5G). None of the top 5 pathways overlapped between cell lines and none of the top 5 pathways in CHP-100 or TTC-466 are shared with the core LSD1 pathways previously defined (Figure 4G). This raises an interesting question regarding the distinct enzymatic and nonenzymatic functions of LSD1 in EwS cell lines. Emerging evidence indicates that LSD1 performs critical scaffolding and recruitment functions independently of its enzymatic activity in other contexts^15,49^. Additionally, the reversible noncompetitive LSD1 inhibitor, SP-2509, was originally theorized to be more cytotoxic than other LSD1 enzymatic inhibitors due to its ability to disrupt the nonenzymatic functions of LSD1^50,51^. However, the specific nonenzymatic functions of LSD1 in EwS remains unclear.

**Figure 5:**
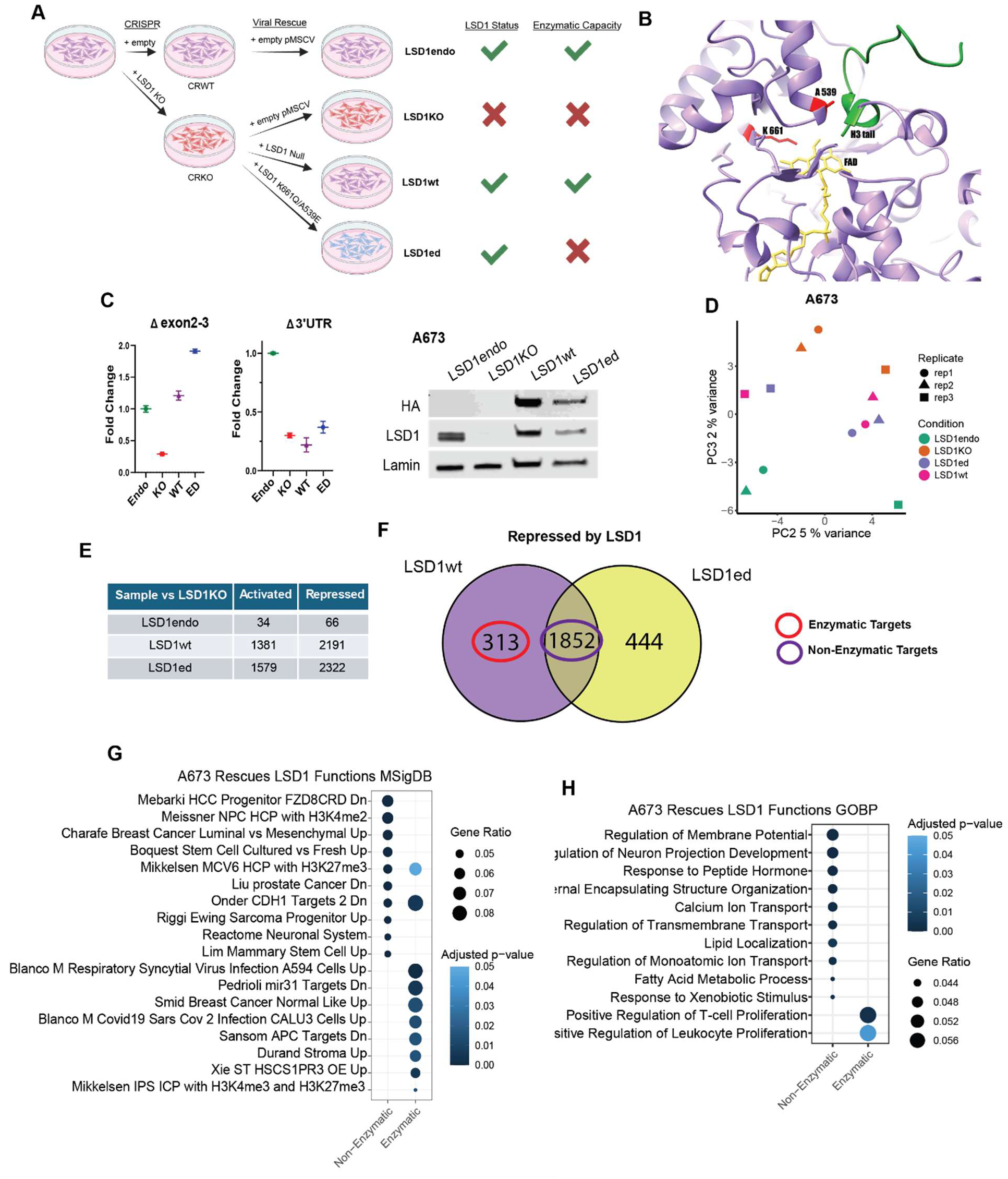
LSD1 functions predominantly nonenzymatically in A673. **(A)** Schematic of KO and viral transduction of LSD1 full length wild type (LSD1wt) and enzymatically dead LSD1 (LSD1ed) harboring K661Q and A539E mutations. **(B)** AlphaFold structure prediction of the enzymatic pocket of LSD1 with K661Q and A539E mutations highlighted. **(C)** Western blot validating rescue of LSD1 protein with the proper 2xHA-tag, and qRT-PCR fold change of regions spanning the exon 2-3 junction and within the 3’UTR. **(D)** Principal component analysis (PCA) of A673 rescue samples comparing PC2 vs PC3. **(E)** Table of genes differentially expressed between the endogenous LSD1 (LSD1endo), LSD1wt, and LSD1ed compared to LSD1KO. **(F)** Overlap of genes downregulated by expression of LSD1wt and LSD1ed. Genes common between LSD1wt and LSD1ed are defined to be nonenzymatic gene targets and are circled in purple. Genes down regulated by LSD1wt but unaffected by LSD1ed are defined to be enzymatically regulated genes and are circled in red. Pathway analysis on both enzymatic and non-enzymatic gene sets ranked by gene ratio in A673 based on **(G)** MSigDB Curated and **(H)** GOBP databases.

### LSD1 functions predominantly nonenzymatically in A673

Having defined common and cell line specific LSD1 target genes, we next sought to use both genetic and pharmacological approaches, as above, to differentiate between LSD1 enzymatic and nonenzymatic functions in EwS cell lines. To our knowledge LSD1 had not yet been knocked down and rescued in an EwS cell line. With our CRISPR KO cell lines, we set out to rescue LSD1 with a HA-tagged full-length wildtype construct (LSD1wt) and a HA-tagged enzymatically dead construct (LSD1ed) delivered by retroviral transduction (Figure 5A). To render LSD1 enzymatically dead, we mutated the lysine at residue 661 to glutamine (K661Q) and the alanine at residue 539 to glutamate (A539E) which interferes with the interaction between the FAD cofactor and the H3 tail, and has been shown to disrupt LSD1 enzymatic activity both on naked DNA and nucleosomes^52^ (Figure 5B). Endogenously, LSD1 has 2 splice isoforms expressed in EwS with and without an additional 60 base pair insert between exons 2 and 3 (LSD1, LSD1 + 2A), but we were unable to doubly rescue both isoforms at the same time. We achieved rescue of LSD1wt and LSD1ed without the 2a exon in both A673 and CHP-100, but not TTC-466, validated by protein and qRT-PCR (Figure 5C, Supplementary Figure 10A-B).

We performed RNA sequencing on our KO cells expressing the rescue constructs to define the genes specifically regulated by LSD1 enzymatic and nonenzymatic functions. However, we found virus treatment itself is the strongest driver of variability (Figure 5D, Supplementary Figure 9A-B). In both cell lines, the number of DEGs between the wild type and LSD1 KO cells (CRWT vs CRKO) is significantly greater than the number of DEGs between these same cells after treatment with an empty virus control (LSD1endo vs LSD1KO) (Supplementary Figure 9C-D). This suggests treatment of cells with virus confounds our ability to analyze the rescue experiments (Supplementary Figure 9A-B). In CHP-100, all the virally treated samples clustered together and separately from the samples not treated with virus (CRKO and CRWT) samples in PCA (Supplementary Figure 9B). As a result, the transcriptomic signature of LSD1 expression was almost completely negated by empty virus treatment. Compared to LSD1KO, rescue with LSD1wt resulted in 8 activated and 3 repressed DEGS, and rescue with LSD1ed resulted in 14 activated and 0 repressed DEGs (Supplementary Figure 9D). As such, we were unable to determine differences between enzymatic and nonenzymatic activity of LSD1 in CHP-100 using this cell line model. While our A673 samples were subject to the same confounding factor of viral treatment, we still saw significant transcriptomic changes with rescue of both LSD1wt and LSD1ed compared to LSD1KO allowing us to compare enzymatic versus nonenzymatic activity in this particular model. Rescue with LSD1wt resulted in 1381 activated and 2191 repressed DEGs, and rescue with LSD1ed resulted in 1579 activated and 2322 repressed DEGs (Figure 5E, Supplementary Figure 9B). Using these DEGs, we differentiated between enzymatic and nonenzymatic functions of LSD1 in A673.

We define the non-enzymatic targets to be the 1852 genes repressed by both LSD1wt and LSD1ed, and the enzymatic targets to be the 313 genes that are solely repressed by LSD1wt and not LSD1ed (Figure 5F). There was a similar ratio of genes that are activated by LSD1 non-enzymatically (1039) and enzymatically (310) although there were no enriched pathways activated enzymatically (Supplementary Figure 10C-E). Pathway analysis on LSD1 repressed genes revealed CDH1 target genes to be repressed both enzymatically and nonenzymatically, however, there were more nonenzymatic genes in that pathway (Figure 5G). In addition, EwS progenitor gene targets and the neural reactome are repressed nonenzymatically. Only two pathways from Gene Ontology Biological Processes were enriched by LSD1 enzymatic targets (Figure 5H). Since there are no transcriptomic changes with OG-L002 mediated enzymatic inhibition, our data support LSD1 to function predominantly nonenzymatically in A673 cells.

### Repression of CDH1 target genes is nonenzymatically regulated

Due to the limitations of the genetic rescues, we took an orthogonal pharmacological approach combining KO, UM171 and OG-L002 treated cells to define genes that LSD1 regulates enzymatically and nonenzymatically. Using this method, we define our nonenzymatically regulated genes to be those common between KO and UM171 treatment but not OG-L002 treatment, and our enzymatically regulated genes as those common between all three. In CHP-100 there are 104 nonenzymatically and 127 enzymatically upregulated genes (Figure 6A). In TTC-466 there are 75 nonenzymatically and 34 nonenzymatically upregulated genes (Figure 6B). Using this approach, there are 555 nonenzymatically and 0 enzymatically upregulated DEGs in A673 (Figure 1E, 2D). There are 50 downregulated nonenzymatic DEGs in A673, 14 downregulated nonenzymatic DEGs in CHP-100 and none in TTC-466 (Supplementary Figure 11A-C). We performed pathway analysis on these gene sets in each cell line. In both CHP-100 and TTC-466, repression of CDH1 target genes is regulated nonenzymatically in addition to other EMT pathways (Figure 6C-D). Of note, in CHP-100 LSD1 nonenzymatically co-regulated Ewing sarcoma progenitor targets and in TTC-466, our results align with previous published LSD1 targets (Figure 6C-D). Thus in light of our data showing OG-L002 has no transcriptional effect in A673 cells, repression of CDH1 is regulated nonenzymatically by LSD1 in all three cell lines, suggesting LSD1 represses these gene targets in a different manner than previously shown in epithelial cells^18^. Nevertheless, our results address a longstanding question in the field and define the specific enzymatic and nonenzymatic gene targets of LSD1 in EwS cell lines.

**Figure 6:**
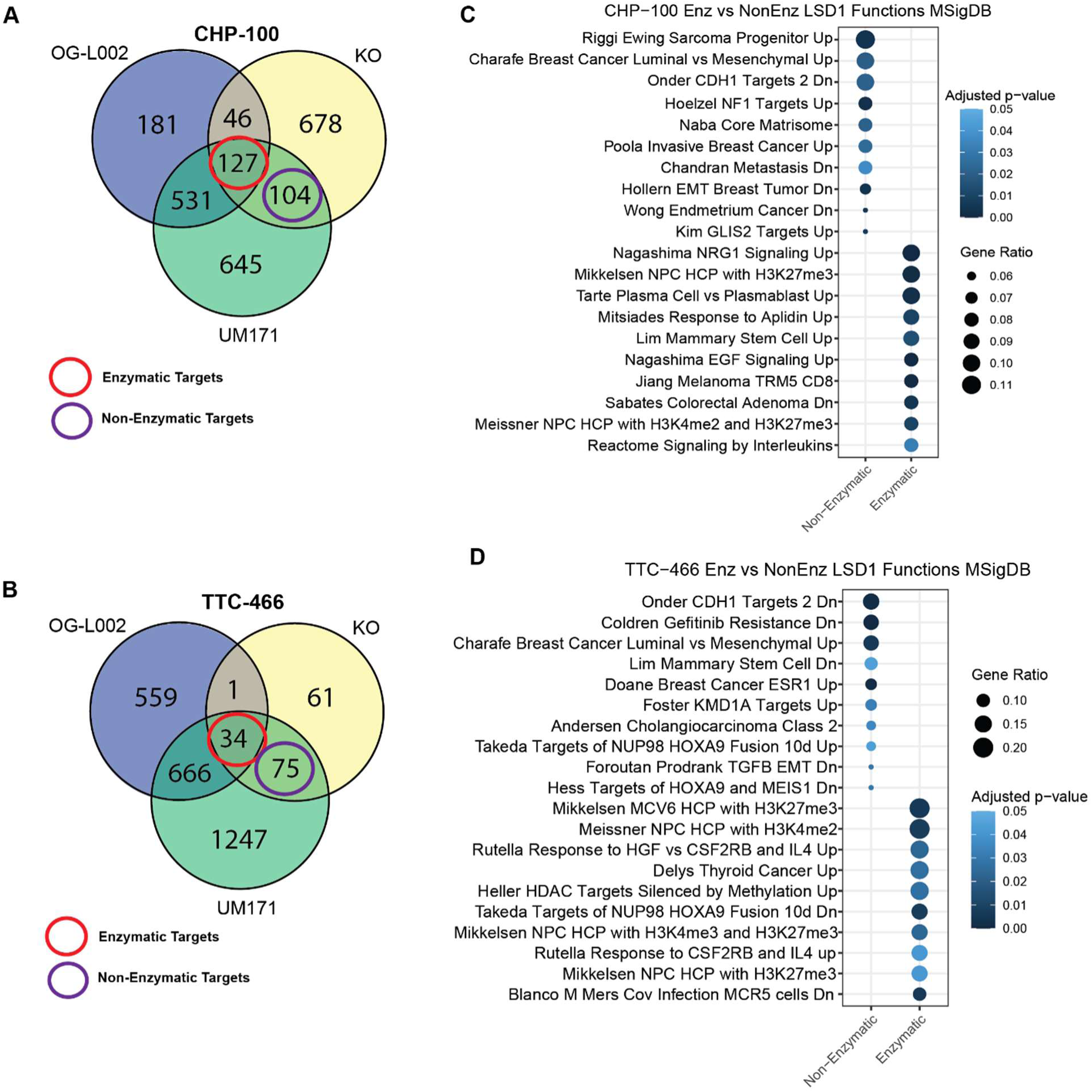
Repression of CDH1 target genes is nonenzymatically regulated. Overlap of genes differentially upregulated by LSD1 KO, UM171 treatment, and OG-L002 treatment in **(A)** CHP-100 and **(B)** TTC-466. Genes commonly up regulated by all three conditions are defined to be enzymatic targets and are circled in red. Genes that are upregulated by KO and UM171 but are not affected by OG-L002 are defined to be nonenzymatic targets and are circled in purple. Pathway analysis on both gene sets in **(C)** CHP-100 and **(D)** TTC-466 ranked by gene ratio from the MSigDB Curated data base.

## DISCUSSION

There has been considerable interest in LSD1 inhibition in EwS since the early 2010s when it was discovered to be overexpressed in EwS^24^. Since then, LSD1 has been shown to correlate with poor patient prognosis, to co-localize with EWSR1::FLI1, and to regulate a similar gene set as EWSR1::FLI1^25,29^. However, we recently demonstrated that the off target pharmacological effects of noncompetitive LSD1 inhibition are responsible for the disruption of the fusion transcriptional program^30^. Therefore, the specific function of LSD1 in EwS remained undefined, both enzymatically and nonenzymatically. Through a combination of genetic and pharmacological techniques, we identified gene sets commonly regulated by LSD1 across three cell lines, including repression of neurotransmitter synaptic functioning and e-cadherin target genes. Of note, these pathways were predominantly regulated by non-canonical nonenzymatic functions of LSD1 (Figure 7). Through use of multiple orthogonal depletion methods and analysis of proliferation, colony formation, and gene expression in a small panel of EwS cell lines, we uncovered the potential of 3D growth assays to identify cell line-specific sensitivity to LSD1 that hasn’t been explored.

**Figure 7:**
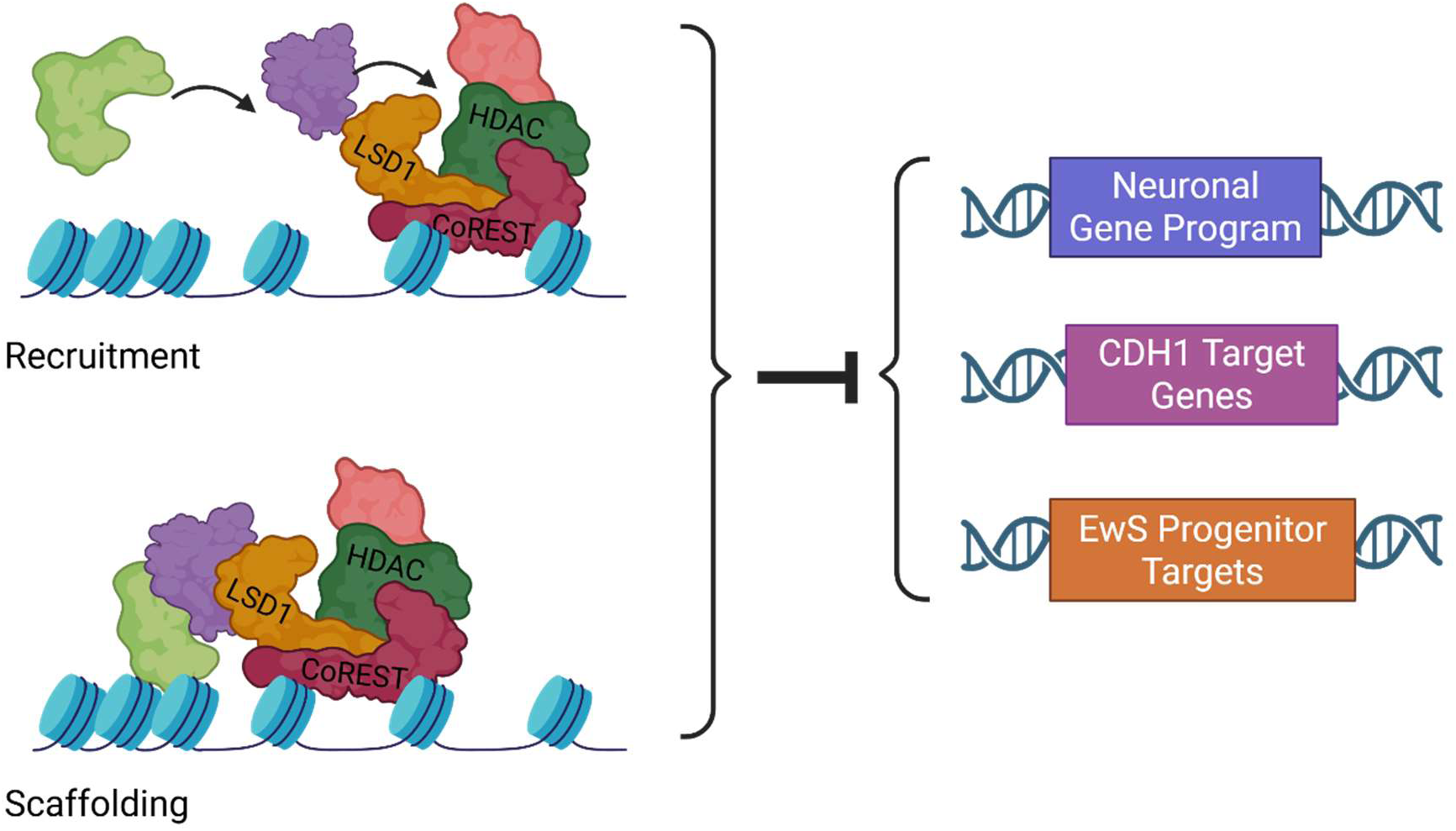
**Model figure depicting nonenzymatic roles of LSD1 in Ewing sarcoma that contribute to the oncogenic program of these cell lines. Image created in BioRender**.

The power of our study comes from the use of orthogonal genetic and pharmacological methods to define our LSD1 regulated gene sets. However, there are some limitations of our approach. shRNA mediated knock down is a longstanding method of choice to study protein function, yet our results suggest that viral treatment of cells was the main driver of phenotypic changes seen with LSD1i, and thus we were unable to use this technique. Cells undergoing CRIPSR-Cas9 gene editing are placed under a significant selective pressure during electroporation that selects for cells with the strongest growth potential, changing the phenotype of the cells and possibly masking a signal due to LSD1 loss^30^. UM171 directly binds HDACs and CoREST, in addition to LSD1, so this degrader is not LSD1 specific^42,43^. Furthermore, use of viral transduction to express LSD1 constructs significantly masked the transcriptomic changes resulting from LSD1 expression. Despite these limitations, we leveraged a range of available techniques in parallel to curate a robust data set defining LSD1 gene regulation in EwS.

Our transcriptomic studies revealed the role of LSD1 in repressing neurotransmitter synaptic functioning. These are pathways that are activated in neurological development, and since LSD1 is highly expressed in early development, our data suggests LSD1 is repressing neurological development in EwS. This finding aligns with the established role that LSD1 represses neurological differentiation in non-neuronal tissues^53^. However, there is also debate surrounding the cell of origin for EwS whether it derives from a neural crest stem cell or a mesenchymal stem cell. Results from Vasileva et al demonstrate that neural crest stem cells can express EWSR1::FLI1 leading to tumors in zebrafish^54^. Furthermore, data from Miller et. al show EwS cells exist somewhere along an axis of neuroendocrine to mesenchymal and pluripotent to differentiated^55,56^. In this case, LSD1 could be contributing to holding cells in a more neurological progenitor, undifferentiated state, and therefore in a more aggressive state. This role of LSD1 is consistent with patient data demonstrating higher levels of LSD1 correlate to poor patient prognosis^25^.

Further supporting LSD1’s contribution to a more aggressive phenotype is the role we uncovered for LSD1 repressing e-cadherin (CDH1) target genes. This creates a pseudo “loss of e-cadherin” environment which is a hallmark of epithelial-to-mesenchymal transition and metastasis. There is prior evidence that LSD1 is recruited through the SNAG domain of Snai1 to repress e-cadherin by demethylating H3K4me2 at the e-cadherin promoter^18^. Our data suggests LSD1 represses CDH1 target genes in a demethylase-independent mechanism, possibly by HDAC recruitment or another scaffolding role. An alternative explanation could be LSD1 interacting with a metastasis associated (MTA) protein as a member of the NuRD complex. However, previous reports have shown this interaction is anti-metastatic in breast cancer. Our results thus support a model where LSD1 represses CHD1 targets nonenzymatically in EwS contributing to a pro-metastatic environment.

Considering these results, we were surprised to see that while loss of LSD1 does not consistently impact 2D cellular proliferation, there are cell line specific defects in 3D growth when treated with UM171. We first rationalized this finding to be the result of off targets of UM171. However, we see the same cell line dependent response when we treat with enzymatic inhibitor, OG-L002, which binds tightly and specifically to LSD1. Instead, 2D toxicity or proliferation assays may be insufficient as a reporter assay for LSD1 function. The only 3D growth assay of TCP derivatives published to date solely included A673 cells, missing potential effects in other cell lines^48^. It is reasonable that LSD1’s role in repressing e-cadherin target genes contributes to the observed defect in 3D growth by reducing cell-to-cell junctions. While we do not consistently see a 3D growth delay in all cell lines, A673 is a particularly odd cell line as it harbors a BRAF mutation and is one of the only EwS cell lines able to tolerate fusion knock down. Another possible explanation is that OG-L002 has been shown to disrupt LSD1 binding to SNAG domain proteins. Differing expression of SNAG domain containing proteins in different cell lines could account for the varied response to OG-L002. Future experiments using new LSD1 enzymatic inhibitors with Grob fragmentation that do not disrupt SNAG binding are needed to ascertain whether SNAG domain containing proteins account for the cell line specific response to LSD1 inhibition or whether this 3D growth defect is strictly due to enzymatic disruption.

Taken together, our data begins to answer longstanding questions surrounding the function of LSD1 in Ewing sarcoma and supports a small and distinct role for LSD1 mediated gene regulation in Ewing sarcoma. We set out to define the enzymatic vs nonenzymatic LSD1 functions in Ewing sarcoma and have uncovered a previously undefined role of LSD1 in repressing neurotransmitter functioning at synapses and CDH1 target genes. Both roles could promote a more developmental and metastatic environment, contributing to the worse outcomes seen for patients with higher LSD1 expression.

Furthermore, LSD1 predominantly functions nonenzymatically in these pathways. With a new emphasis on noncanonical functions of LSD1 it is even more crucial to characterize the full scope of LSD1 complexes to define the mechanism by which LSD1 regulates these genes. Our data suggests that certain cell lines, and by extension certain tumors, could be susceptible to LSD1 inhibition targeting these predominant nonenzymatic functions and supporting an avenue where LSD1 inhibitor therapy is indicated. LSD1 inhibitors are currently in clinical trials for other cancers and could be repurposed and used as a combinational therapeutic for certain LSD1 sensitive Ewing sarcoma tumors.

## ACKNOWLEDGEMENTS

We thank the High-Performance Computing group at Nationwide Children’s Hospital for their support; the Institute for Genomic Medicine at Nationwide Children’s Hospital for sequencing support: and the CRIPSR Core at Nationwide Children’s Hospital for reagent generation support. We are grateful to the members of the Theisen Lab, including Michelle Cruz, for comments and discussion of this manuscript during its preparation. This research was supported by institutional startup funds awarded to E.R.T., the American Cancer Society Research Scholar Grant RSG-22-118-01-DMC awarded to E.R.T., a St. Baldrick’s Scholar Award to E.R.T., a The Ohio State University Presidential Fellowship to R.D.D., a The Ohio State University Alumni Grants for Graduate Research and Scholarship Grant to R.D.D., a CancerFree KIDS New Idea Award to R.D.D., and NIH T32 CA269052 for both A.B and J.W.S.

