## Supplementary Materials for "LSD1 Performs Demethylase-Independent and Context-Specific Roles in Ewing Sarcoma"

### SUPPLEMENTARY FIGURES

**Supplementary Figure 1:** Western blot images of all replicates in each cell line (A673, CHP-100, and TTC-466) of **(A)** LSD1 KO **(B)** UM171 degradation with 800nM for 96 hrs, and **(C)** shRNA LSD1 knockdown with 72 hrs of puromycin selection. 2D cellular proliferation and 3D proliferation in agars with representative plates and the corresponding quantification of colonies across replicates in **(D)** A673, **(E)** CHP-100, and **(F)** TTC-466. **(G)** Principal component analysis (PCA) of A673 samples with CRISPR KO, UM171 degradation time points and shRNA LSD1 knockdown. **(H)** PCA analysis of all samples across the three cell lines.

**Supplementary Figure 2.** PCA of KO and UM samples with corresponding Jaccard similarity index, calculated by fisher's exact test, for up- and downregulated genes in **(A)** A673, **(B)** CHP-100, and **(C)** TTC-466.

**Supplementary Figure 3.** Pathway analysis ranked by gene ratio on the overlap of KO and UM171 upregulated genes from MSigDB Curated and GOBP databases in **(A-B)** A673, **(C-D)** CHP-100, and **(E-F)** TTC-466.

**Supplementary Figure 4.** Commonly **(A)** down- and **(B)** upregulated genes between the fusion of all three cell lines, *EWSR1::FLI1* type 1 in A673, *EWSR1::FLI1* type 2 in CHP-100 and *EWSR1::ERG* in TTC-466. **(C)** Overlap of genes differentially up- and downregulated common between core fusion targets and core LSD1 repressed targets between all three cell lines. Genes commonly up and downregulated by the fusion overlapped with genes that are upregulated by LSD1 loss, as defined by the overlap of KO and UM171 treatment, in **(D)** A673, **(E)** CHP-100, and **(F)** TTC-466. All cell line specific overlaps (D-F) have a p-values < 0.001, determined by chi-square test. Gene set enrichment analysis of genes upregulated by LSD1 loss as the gene set compared to a rank order list of genes differentially expressed by the rescue of the fusion compared to KO of the fusion in **(G)** A673, **(H)** CHP-100, and **(I)** TTC-466. **(J)** Western blot validation and of CHP-100 knockdown and rescue of *EWSR1::FLI1*. **(K)** Validation of loss and rescue of 3D colony growth in agar with knock down and rescue of *EWSR1::FLI1* in CHP-100. Validation of A673 and TTC-466 have been previously published[29,57]. **(L)** Quantification of colonies in CHP-100 with KD/rescue of *EWSR1::FLI1.*

**Supplementary Figure 5. (A)** Western blot demonstrating LSD1 depletion over time from 0.5-6hrs of 800nM of UM171 treatment. **(B)** Western blots of all replicates of cells treated with 800nM of UM171 for 6 and 24hrs in all three cell lines compared to DMSO. Four-way overlap of genes differentially up- and downregulated between KO vs WT, and 6, 24, and 96hrs of 800nM of UM171 vs DMSO in **(C)** A673, **(D)** CHP-100, and **(E)** TTC-466. Overlap of genes commonly upregulated between KO and UM171 in all three cell lines at **(F)** all 3 UM171 time points**, (G)** 6hrs of UM171, and **(H)** 24hrs of UM171.

**Supplementary Figure 6.** Pathway analysis ranked by gene ratio from MSigDB Curated and GOBP databases of the genes commonly upregulated between KO and UM171 at 6, 24, and 96hrs of treatment in **(A-B)** CHP-100 and **(C-D)** TTC-466. There are no enriched pathways in either cell line for the overlap of KO and 6hr UM171 in GOBP.

**Supplementary Figure 7.** 2D cellular proliferation over 1 week and 3D growth in soft agar over 2 weeks with a representative plate and the corresponding quantification of colonies across replicates in **(A)** A673, **(B)** CHP-100, and **(C)** TTC-466 of 800nM of UM171 for 6hrs compared to DMSO. **(D)** Dose response curve from 96hrs of UM171 treatment in each cell line. **(E)** RNA and **(F)** protein levels of LSD1 with DMSO and OG-L002 treatment in each cell line.

**Supplementary Figure 8.** PCAs of all cell lines treated with 2µM of OG-L002 vs DMSO comparing PC1 Vs PC2 **(A)** and PC2vs PC3 **(B)**.

**Supplementary Figure 9.** A673 and CHP-100 PCAs of **(A)** LSD1endo, LSD1KO, LSD1wt, and LSD1ed rescue constructs and **(B)** CRISPR LSD1 KO and LSD1 WT cells. **(C-D)** Tables of genes differentially expressed in A673 and CHP-100 between CRISPR KO vs WT, empty vector treated LSD1KO vs

LSD1endo, LSD1wt, and LSD1ed, CRISPR WT vs empty vector treated LSD1endo, and CRISPR KO vs empty vector treated LSD1KO.

**Supplementary Figure 10. (A)** Western blot validation of CHP-100 properly 2xHA tagged protein rescue of LSD1wt and LSD1ed. **(B)** qrtPCR fold change of regions spanning the exon 2-3 junction and within the 3'UTR. **(C)** Overlap of genes downregulated by expression of LSD1wt and LSD1ed. Genes common between LSD1wt and LSD1ed are defined to be nonenzymatic gene targets and are circled in purple. Genes down regulated by LSD1wt but unaffected by LSD1ed are defined to be enzymatically regulated genes and are circled in red. Pathway analysis on both enzymatic and non-enzymatic gene sets ranked by gene ratio in A673 based on **(D)** MSigDB Curated and **(E)** GOBP databases.

**Supplementary Figure 11.** Overlap of genes differentially downregulated by LSD1 KO, UM171 treatment, and OG-L002 treatment in **(A)** CHP-100, **(B)** TTC-466, and **(C)** A673. Genes commonly downregulated by all three conditions are defined to be enzymatic targets and are circled in red. Genes that are downregulated by KO and UM171 but are not affected by OG-L002 are defined to be nonenzymatic targets and are circled in purple.

**SUPPLEMENTARY TABLE**

**Supplementary Table 1.** Sequences of LSD1 cloning PCR primers, qrtPCR primers, LSD1 wild type and K661Q/A539E rescue construct gene blocks, and CRISPR KO guides.

Supplementary Figure 1

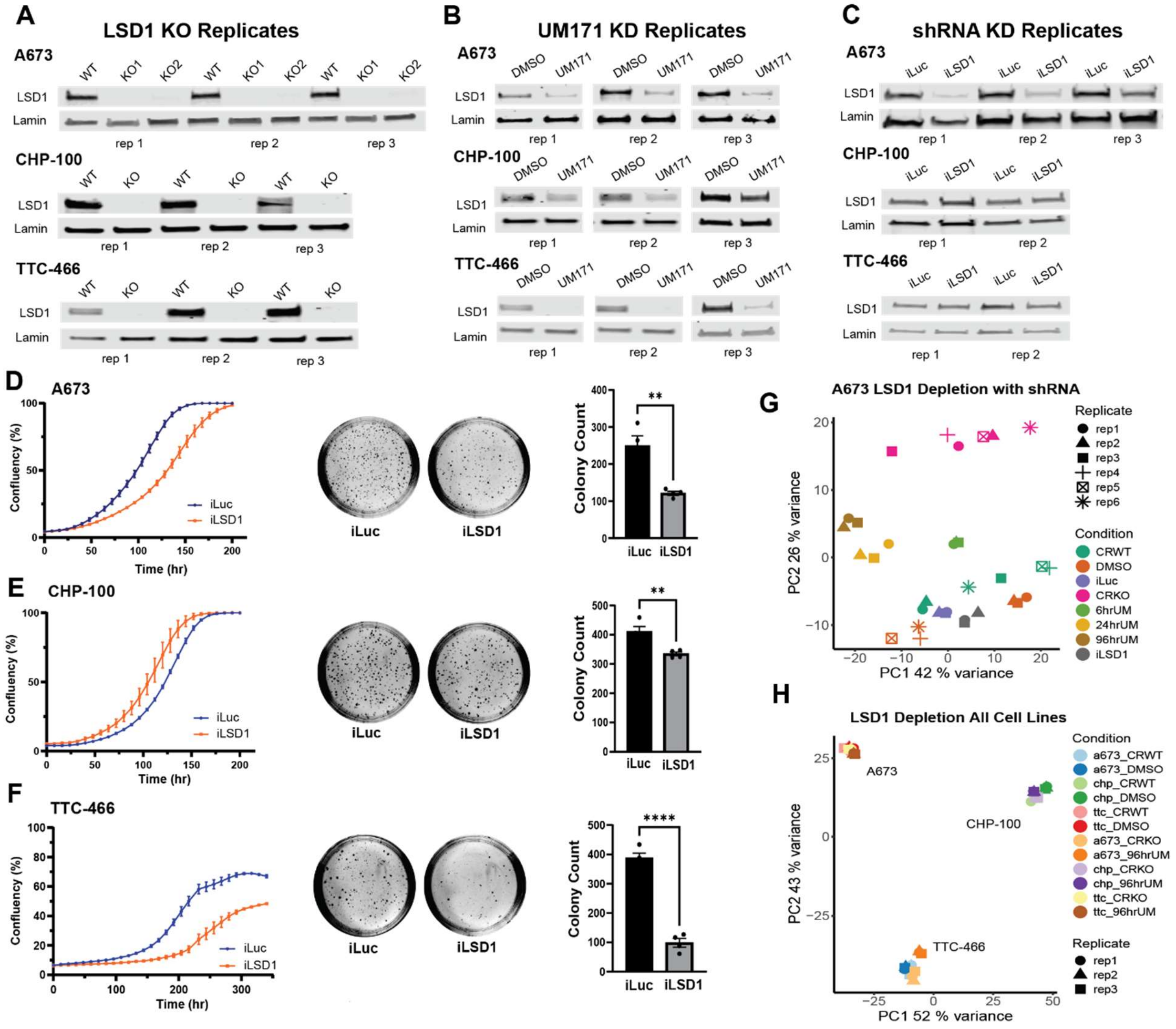

Supplementary Figure 2

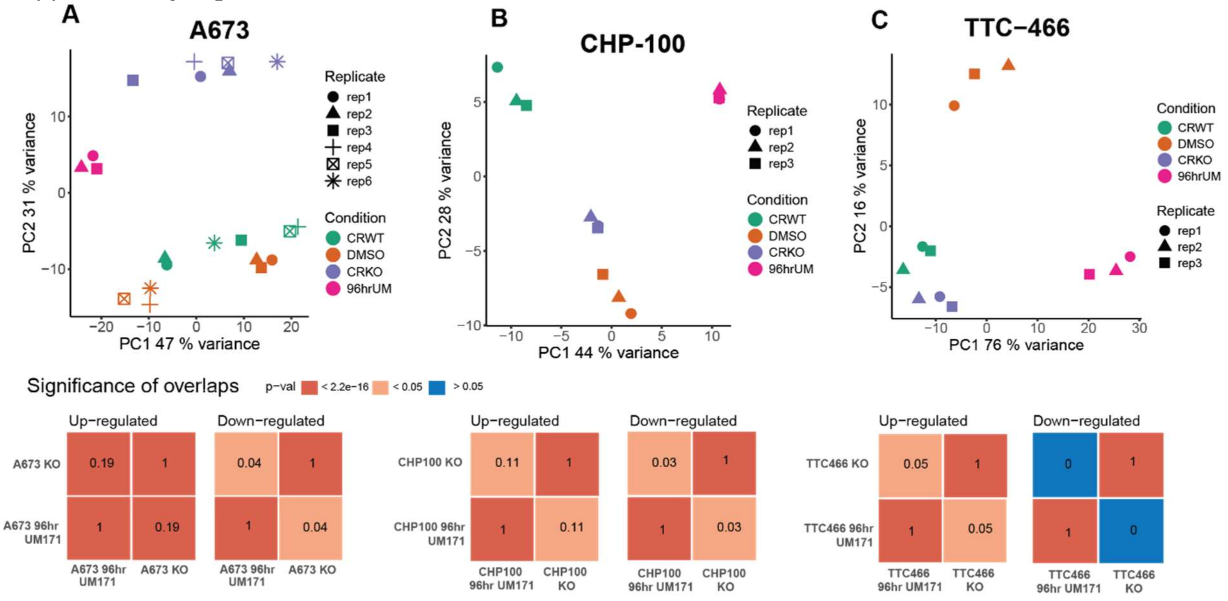

Supplementary Figure 3

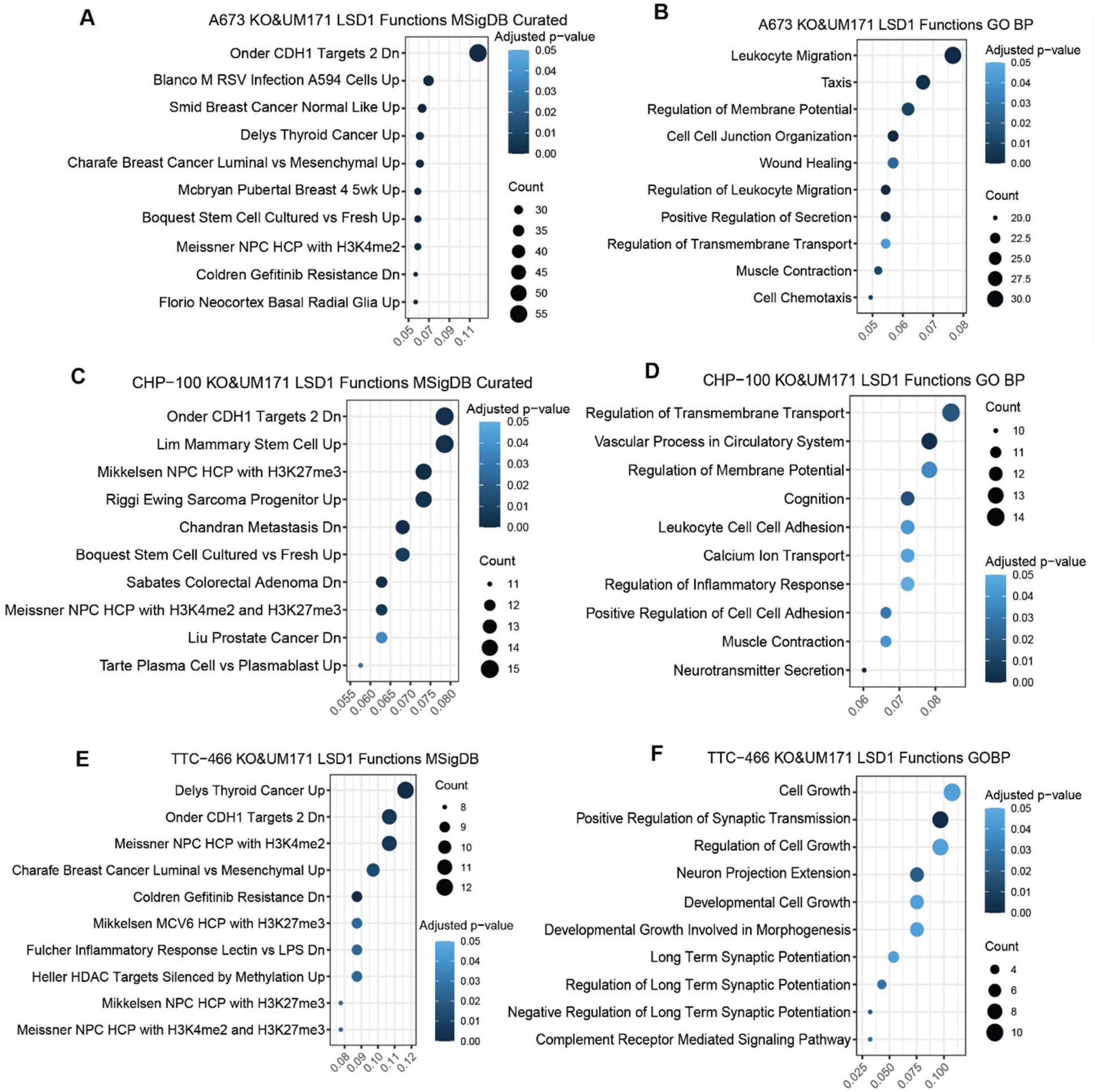

Supplementary Figure 4

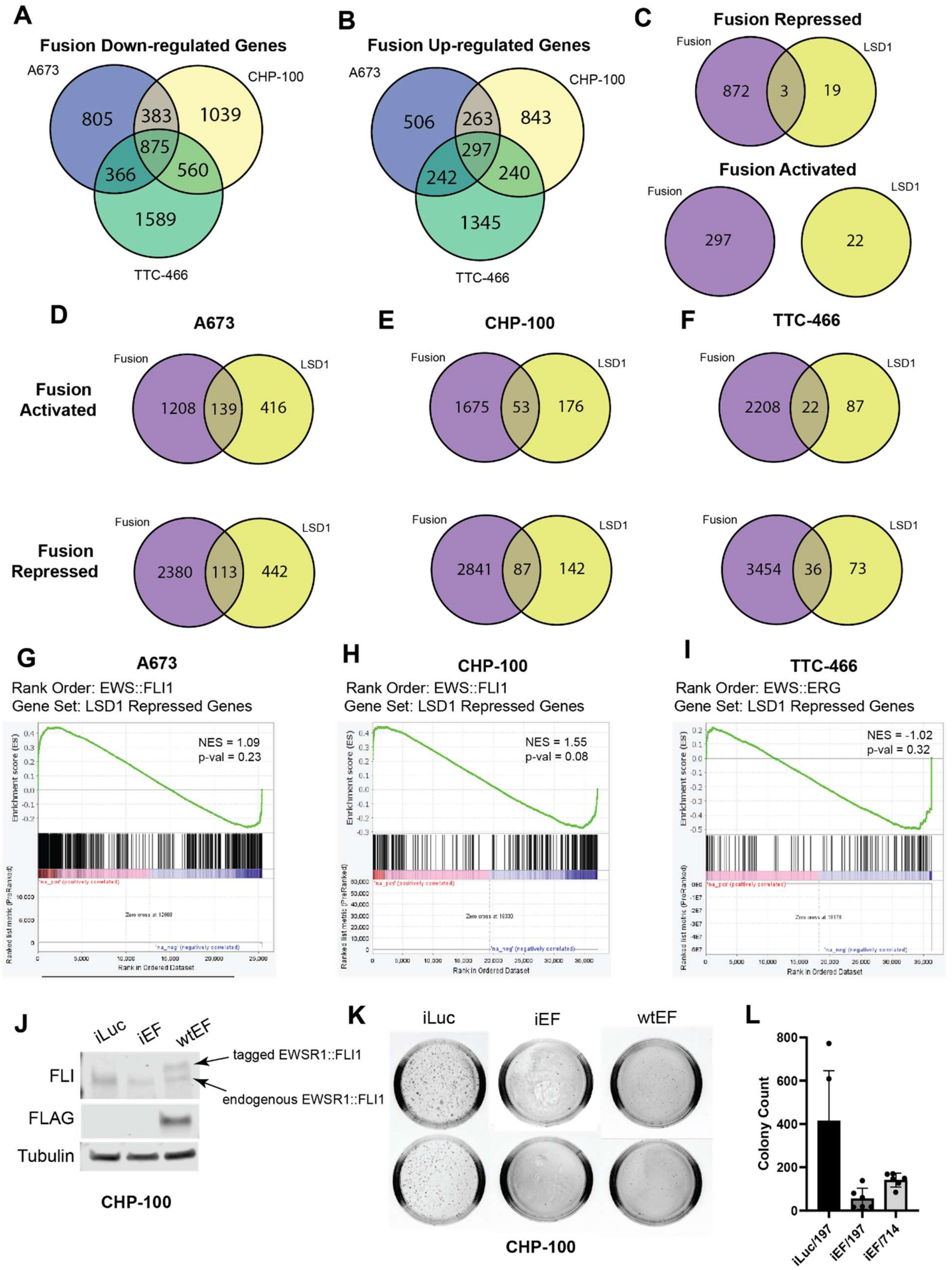

Supplementary Figure 5

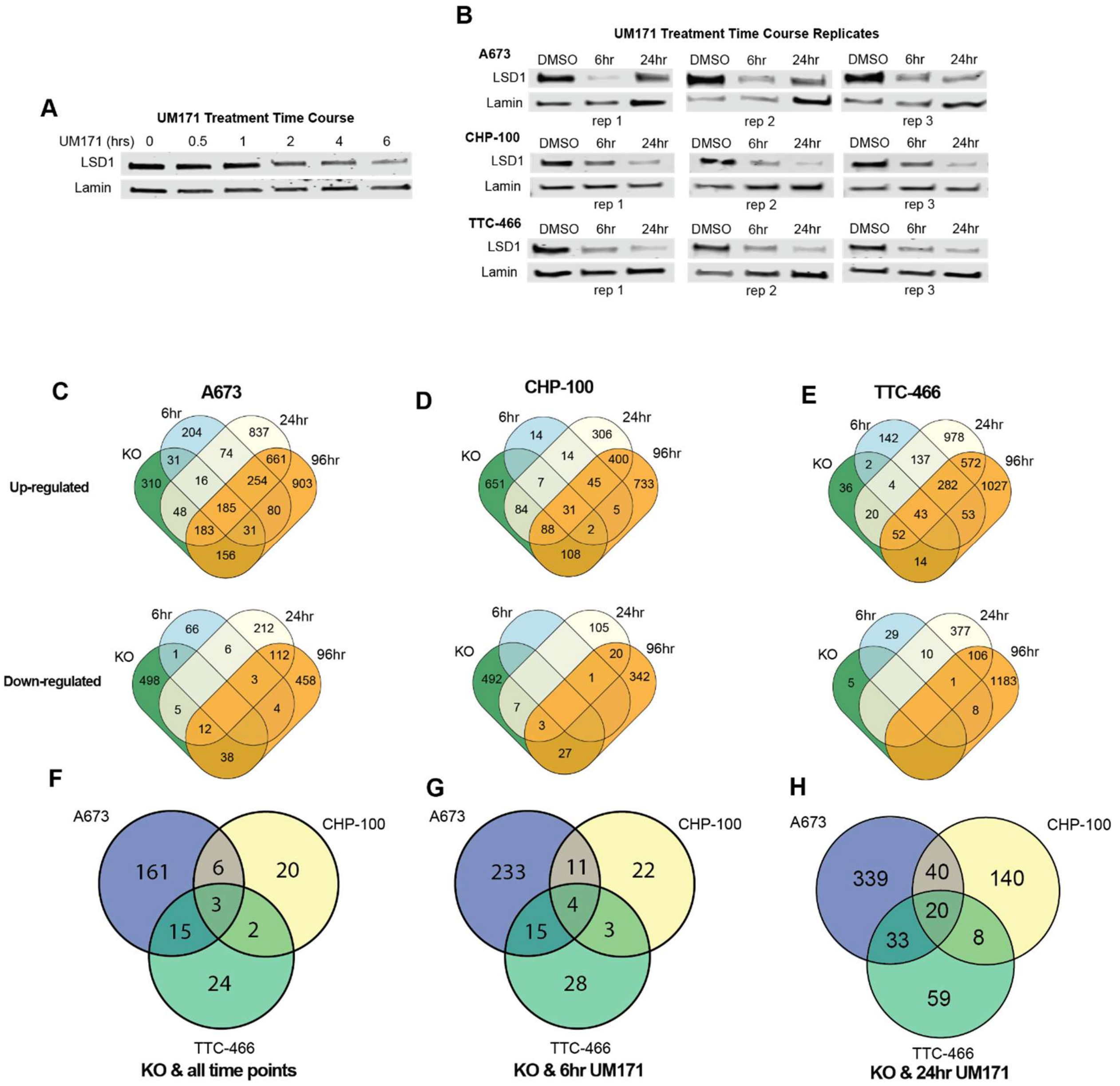

Supplementary Figure 6

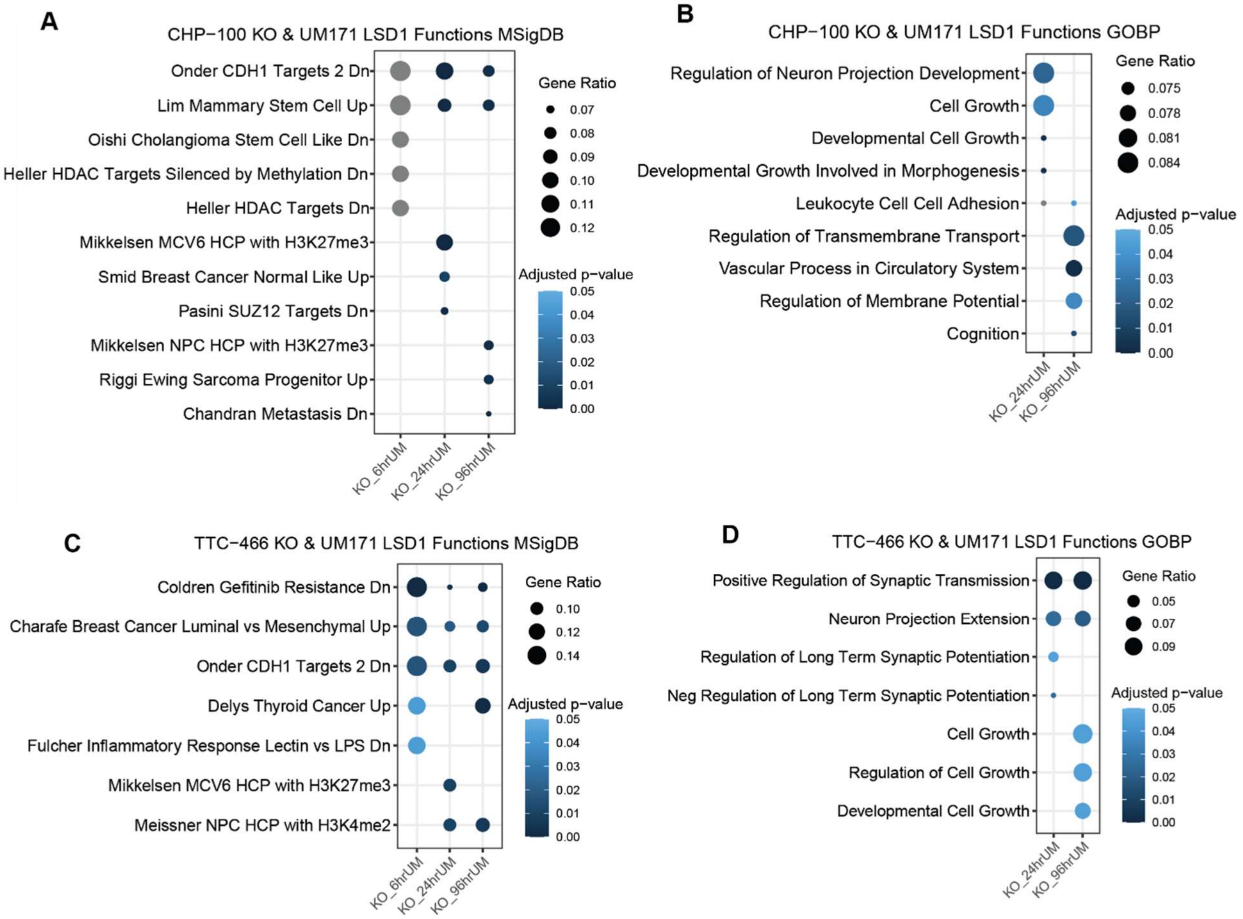

83 Supplementary Figure 7

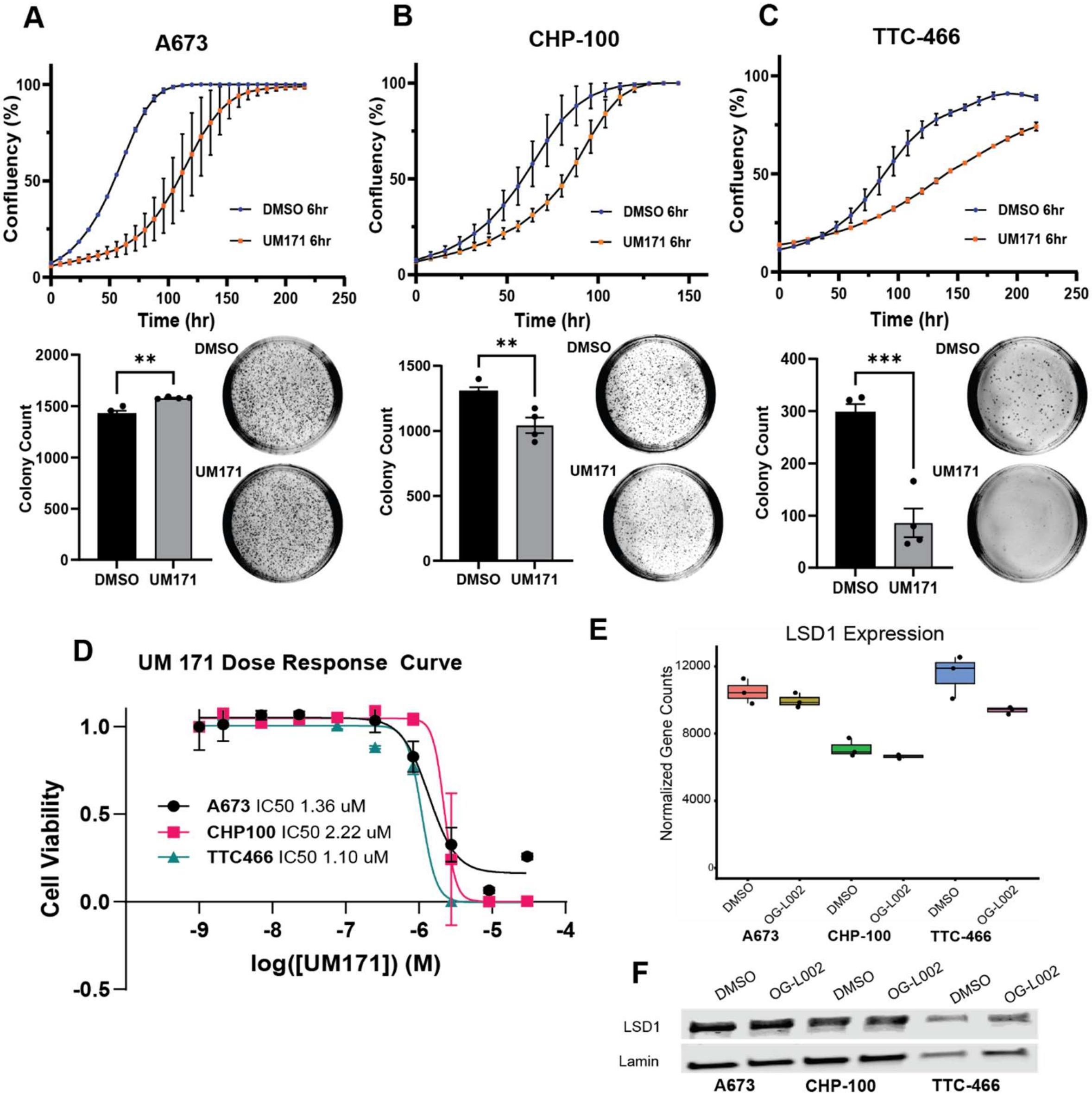

84

Supplementary Figure 8

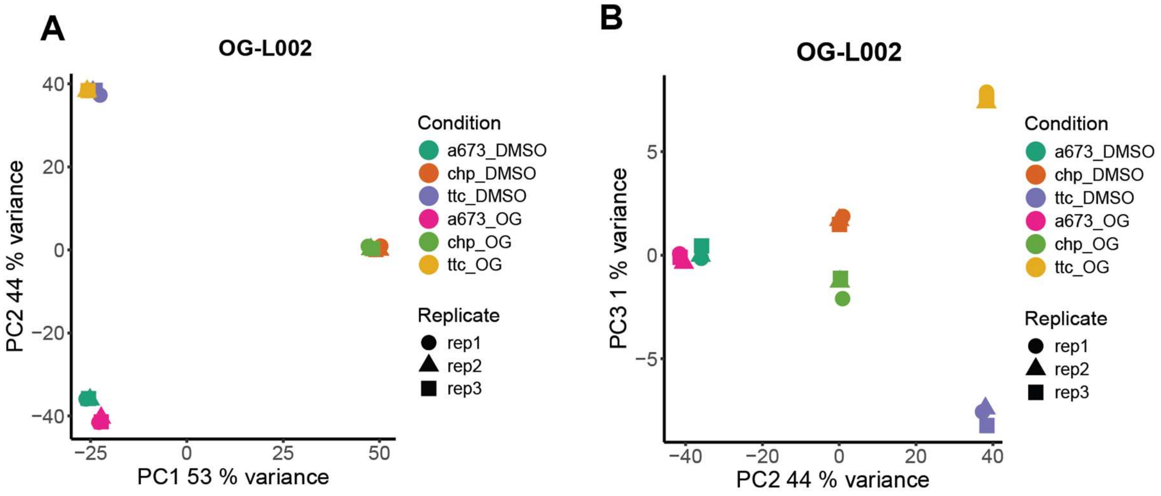

Supplementary Figure 9

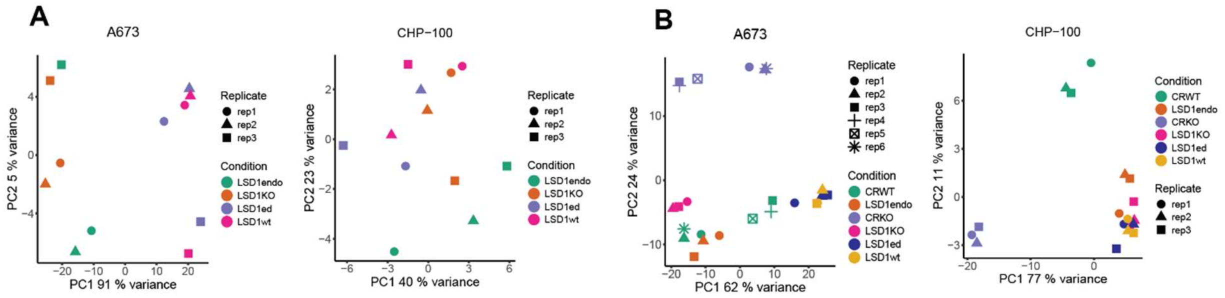

C

| A673 Comparison | Activated | Repressed |
|---|---|---|
| CRWT vs CRKO | 554 | 960 |
| LSD1endo vs LSD1KO | 34 | 66 |
| LSD1wt vs LSD1KO | 1381 | 2191 |
| LSD1ed vs LSD1KO | 1579 | 2322 |
| LSD1endo vs CRWT | 86 | 78 |
| LSD1KO vs CRKO | 1074 | 937 |

D

| CHP-100 Comparison | Activated | Repressed |
|---|---|---|
| CRWT vs CRKO | 529 | 971 |
| LSD1endo vs LSD1KO | 31 | 8 |
| LSD1wt vs LSD1KO | 8 | 3 |
| LSD1ed vs LSD1KO | 14 | 0 |
| LSD1endo vs CRWT | 264 | 196 |
| LSD1KO vs CRKO | 756 | 1514 |

90 Supplementary Figure 10

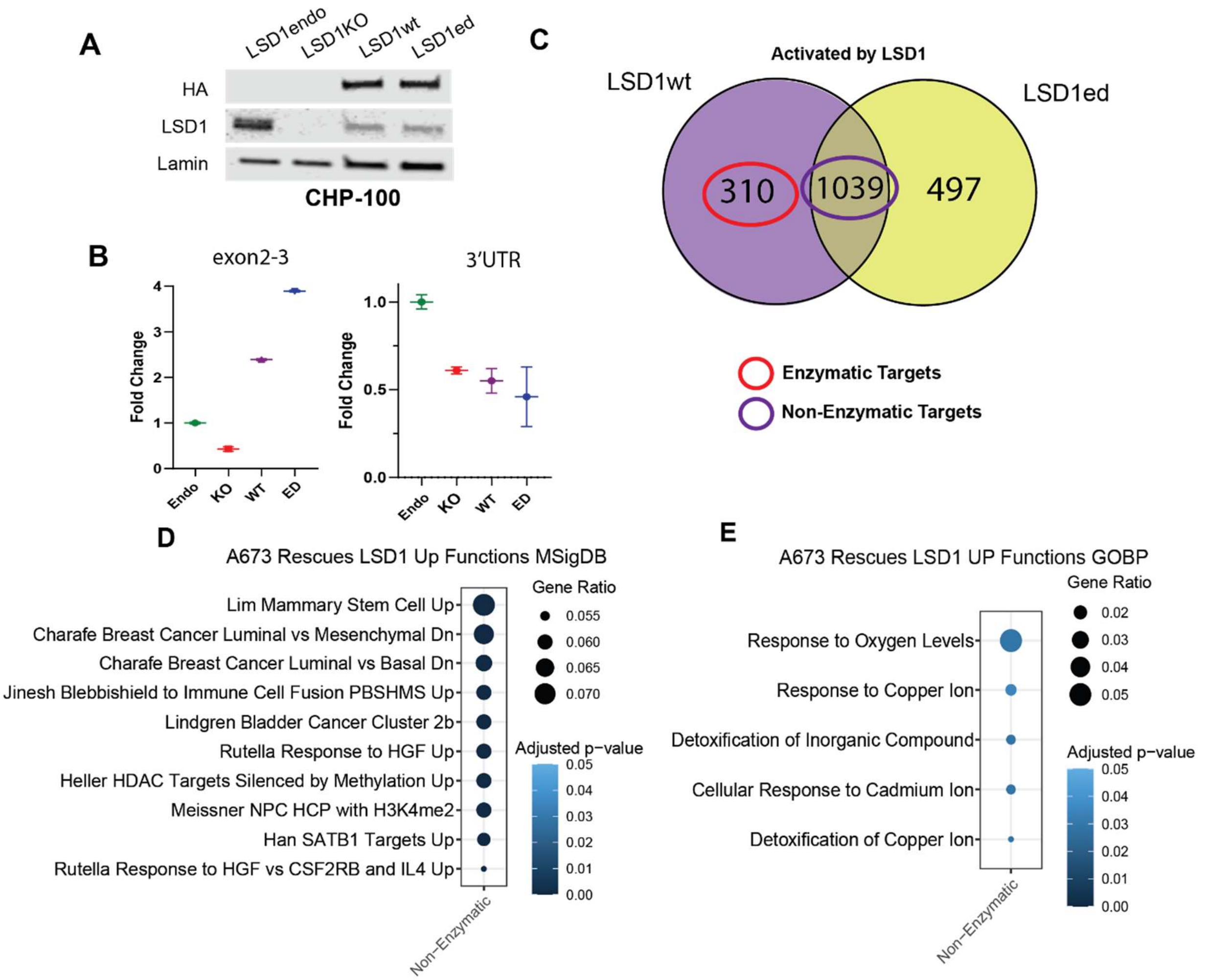

91

Supplementary Figure 11

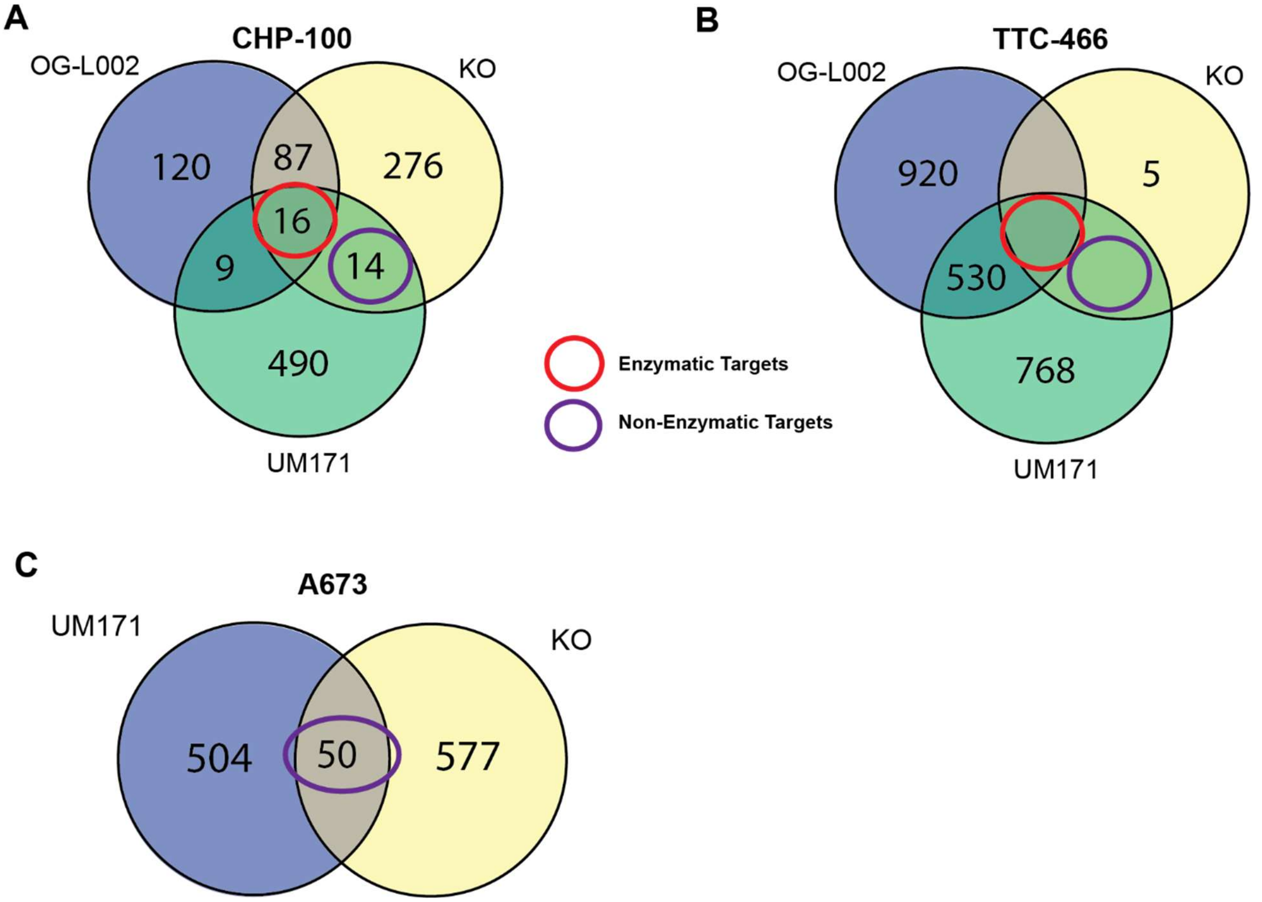

| **Supplementary Table 1** | |
|---|---|
| **LSD1 PCR Primers** | **Sequence** |
| LSD1_for_gibson | TTATCTGGGAAGAAGGCGGCAGCC |
| LSD1_rev_gibson | GGTAGAATTCGTTAACCTCACATGCTTGGGGACTGC |
| **qRTPCR primers** | **Sequence** |
| LSD1KO_exon2_3_F | CCTCAAGCCCCACCTGAGGA |
| LSD1KO_exon2_3_R | CTGGGTCTGTTGTGGTCCACTG |
| LSD1KO_3UTR_F | GCCCATGTGCCTGTTTCTGCC |
| LSD1KO_3UTR_R | GACAAGTTCCTCCCTGTGCTCTAGG |
| RPL30_F | GGGGTACAAGCAGACTCTGAAG |
| RPL30_R | ATGGACACCAGTTTTAGCCAAC |
| **Construct Gene blocks** | **Sequence** |
| TSHA_LSD1 | AGGCGCCGGAATTAGATCTCATGAGTGCTTGGAGTCATCCACAATTCGAAAAAG<br>GTGGAGGTTCCGGAGGTGGATCGGGAGGTTCTGCCTGGTCTCATCCACAATTTG<br>AGAAGGGCGGCTACCCATACGATGTTCCAGATTACGCTTACCCATACGATGTTCC<br>AGATTACGCTGGCTCCTTATCTGGGAAGAAGGCGGCAGCCGCGGCGGCGGCGG<br>CTG |
| TSHA_LSD1_K661Q_A539E | GCGAATCCCCCAAGTGATGTATATCTCTCATCAAGAGACAGACAAATACTTGATT<br>GGCATTTTGCAAATCTTGAATTTGAGAATGCCACACCTCTCTCAACTCTCTCCCTT<br>AAGCACTGGGATCAGGATGATGACTTTGAGTTCACTGGCAGCCACCTGACAGTA<br>AGGAATGGCTACTCGTGTGTGCCTGTGGCTTTAGCAGAAGGCCTAGACATTAAA<br>CTGAATACAGCAGTGCGACAGGTTCGCTACACGGCTTCAGGATGTGAAGTGATA<br>GCTGTGAATACCCGCTCCACGAGTCAAACCTTTATTTATAAATGCGACGCAGTTC<br>TCTGTACCCTTCCCCTGGGTGTGCTGAAGCAGCAGCCACCAGCCGTTCAGTTTGT<br>GCCACCTCTCCCTGAGTGGAAAACATCTGCAGTCCAAAGGATGGGATTTGGCAA<br>CCTTAACCAGGTGGTGTTGTGTTTTGATCGGGTGTTCTGGGATCCAAGTGTCAAT<br>TTGTT |
| **CRISPR KO guides** | |
| LSD1 Guide 3 | TAAAGGTTTGACTCGTGGAG |
| LSD1 Guide 4 | GTCAATTTGTTCGGGCATGT |
